# Spatially Dependent Tissue Distribution of Thyroid Hormones by Plasma Thyroid Hormone Binding Proteins

**DOI:** 10.1101/2023.12.20.572629

**Authors:** Anish D. Bagga, Brian P. Johnson, Qiang Zhang

**Author notes:** Corresponding author: Qiang Zhang, M.D., Ph.D. Gangarosa Department of Environmental Health Rollins School of Public Health Emory University Atlanta, GA 30322. Disclosure Summary: The authors declare no conflict of interest.

## Abstract

Plasma thyroid hormone (TH) binding proteins (THBPs), including thyroxine-binding globulin (TBG), transthyretin (TTR), and albumin (ALB), carry THs to extrathyroidal sites, where THs are unloaded locally and then taken up via membrane transporters into the tissue proper. The respective roles of THBPs in supplying THs for tissue uptake are not completely understood. To investigate this, we developed a spatial human physiologically based kinetic (PBK) model of THs, which produces several novel findings. **(1)** Contrary to postulations that TTR and/or ALB are the major local T4 contributors, the three THBPs may unload comparable amounts of T4 in *Liver*, a rapidly perfused organ; however, their contributions in slowly perfused tissues follow the order of abundances of T4TBG, T4TTR, and T4ALB. The T3 amounts unloaded from or loaded onto THBPs in a tissue acting as a T3 sink or source respectively follow the order of abundance of T3TBG, T3ALB, and T3TTR regardless of perfusion rate. **(2)** Any THBP alone is sufficient to maintain spatially uniform TH tissue distributions. **(3)** The TH amounts unloaded by each THBP species are spatially dependent and nonlinear in a tissue, with ALB being the dominant contributor near the arterial end but conceding to TBG near the venous end. **(4)** Spatial gradients of TH transporters and metabolic enzymes may modulate these contributions, producing spatially invariant or heterogeneous TH tissue concentrations depending on whether the blood-tissue TH exchange operates in near-equilibrium mode. In summary, our modeling provides novel insights into the differential roles of THBPs in local TH tissue distribution.

**Key Points:**

- Thyroxine-binding globulin (TBG), transthyretin (TTR), and albumin (ALB) are plasma thyroid hormone (TH) binding proteins (THBPs) that carry THs from the thyroid gland to extrathyroidal tissues.
- The respective roles of the 3 THBP species in unloading THs once arriving at a tissue are not completely understood.
- Here we developed a spatial human kinetic model of THs and showed that the three THBPs may unload comparable amounts of thyroxine (T4) in the liver but TBG is dominant in contributing T4 in tissues slowly perfused by blood as well as in contributing triiodothyronine (T3) regardless of the tissue’s perfusion rate.
- The TH amounts unloaded by each THBP species are spatially dependent and nonlinear, with ALB being the dominant contributor near the arterial end but conceding to TBG near the venous end in a tissue.
- Our model provides novel insights into the differential roles of THBPs in local TH tissue distribution.

## Introduction

The circulating thyroxine (T4) and triiodothyronine (T3) levels are robustly regulated to stay within narrow (about 2-3 fold) physiological ranges (Jain 2015, Welsh and Soldin 2016). Deviations of the thyroid hormones (THs) from the reference ranges lead to a variety of adverse health outcomes (Combs et al. 2011, Jabbar et al. 2017, Prezioso et al. 2018, Silva et al. 2018, Yavuz et al. 2019). By maintaining the physiological levels of THs, the hypothalamic-pituitary-thyroid (HPT) axis is the primary mechanism for long-term TH homeostasis. To combat transient perturbations, it requires three major TH binding/distributor proteins (THBPs/THDPs) in the blood: thyroxine-binding globulin (TBG), transthyretin (TTR), and albumin (ALB).

The vast majority of circulating THs are bound to and transported by these THBPs (Janssen and Janssen 2017, McLean et al. 2017), with a small fraction also bound to lipoproteins and members of the serine proteinase inhibitors (serpins) superfamily (Benvenga et al. 1988, Benvenga et al. 2002, Benvenga 2013). The molar abundances of the three THBPs in humans follow the order of ALB>TTR>TBG, with ALB more than a hundred times higher than TTR and TTR more than tens of times higher than TBG (Attwood et al. 1978, Gardner and Scott 1980, Attwood and Atkin 1982, Vatassery et al. 1991). However, the binding affinities of THs for the three THBPs follow the reverse order: TBG > TTR > ALB. The binding affinities of T4 for the three THBPs differ by more than two orders of magnitude, while in the case of T3, the binding affinities differ by at least one or two orders of magnitude (Murata et al. 1985, Yabu et al. 1987, Chang et al. 1999, Richardson 2007, McLean et al. 2017). As a result of the vast differences in the THBP abundances and binding affinities, TBG accounts for ∼75% of both plasma total T4 and total T3 despite its low abundance and having only one binding site, TTR accounts for ∼15% of total T4, but less than 5% of total T3, and ALB accounts for less than 5% of total T4 and ∼20% of total T3 (Schussler 2000, Janssen and Janssen 2017, McLean et al. 2017). Overall, less than 0.03% of plasma T4 is free with the vast majority bound to THBPs, while about 0.3% of plasma T3 is free with the vast majority also bound to THBPs (Mendel 1989, Richardson 2007, McLean et al. 2017). TBG is only about 20% saturated and TTR is less than 1% saturated by THs with the majority of their binding sites unoccupied by THs (Schussler 2000, McLean et al. 2017). THBPs are believed to have several functions. (1) They produce a strong buffer in the plasma where a transient increase or decrease of free T4 and T3 can be quickly dampened. (2) THs stored in THBPs can act as a reservoir in the face of short-term TH deficiency and THBPs extend the half-lives of THs. (3) THBPs ensure uniform distribution of circulating THs through a perfused tissue; in the absence of THBPs, a steep gradient of tissue THs would result (Weisiger et al. 1986, Mendel et al. 1987, Mendel et al. 1988).

While a lot has been learned about THBPs, many questions remain unanswered. THs are taken up by tissues predominantly in free form and mainly via cell membrane transporters including MCT8, MCT10, LAT1, LAT2, OATP1c1 etc. (Pizzagalli et al. 2002, Friesema et al. 2003, Bernal et al. 2015, Zevenbergen et al. 2015, Groeneweg et al. 2020). However, the bulk of these THs that are transported into the tissues must first be unloaded off of THBPs in the tissue capillary. The relative contributions of each THBP species in this regard are not completely established but are important for understanding TH tissue distribution as well as potential species differences in TH kinetics. It has been argued that because of the faster dissociation rate constants for the binding between THs and ALB (i.e., shorter residence time of T4 and T3 on an ALB molecule), ALB contributes the most THs to the perfused tissues (Mendel 1989, Schussler 2000). It is also speculated that TTR may be the one that contributes the most T4 to tissues because of its “Goldilocks” (just-about-right) properties among the three THBPs, i.e., intermediate abundance and binding affinity (Robbins 2002, Richardson 2009, Alshehri et al. 2015). Given that TBG is loaded with over two thirds of total T4 and total T3 in the blood, it is also possible that this high abundance compensates for its tight grip on THs such that it makes comparable or even greater contributions than ALB and TTR. In addition, the blood perfuses different tissues at different flow rates, thus THBPs carrying THs would clear through tissues in different amounts of time, during which free THs are transported into and out of tissue proper at potentially different rates. Complicating the situation further is that TH transporters and metabolic enzymes may be differentially expressed in different tissues and heterogeneously expressed in different parts of the same tissues (Halpern et al. 2017, Goemann et al. 2018). It is unclear how all these factors – the residence time of THs on THBPs, THBP abundance, tissue blood transit time, the rates of blood-tissue TH exchange, as well as local metabolism – interplay to determine the amounts of THs that are unloaded from or loaded onto each THBP species in local tissue blood and eventually exchanged with the tissue proper.

While ingenious experimentation can help clarify many of these questions, quantitative understanding of the thyroid system can also benefit greatly from mathematical modeling (DiStefano and Chang 1971, Oppenheimer and Schwartz 1985, Pilo et al. 1990, Curti and Fresco 1992, Eisenberg et al. 2008, McLanahan et al. 2009, Lumen et al. 2013, Berberich et al. 2018, Handa et al. 2021). A physiologically based kinetic (PBK) model of THs describing sufficient details of the THBP binding events with spatial configurations will help dissect the differential roles of THBPs and provide critical quantitative insights. Here we extended a nonspatial human PBK model of THs we have recently published (Bagga et al. 2023) into a spatial model, and reported several novel findings on local TH tissue delivery and contributions by different THBP species.

## Methods

### 1. The nonspatial human PBK model of THs

The details of the nonspatial human PBK model of THs, including its construction and parameterization, were described in our recently published paper (Bagga et al. 2023). The model was rigorously parameterized based on numerous published studies on THBP binding affinities, association/dissociation rate constants, TH and THBP concentrations, plasma/tissue partitioning, half-lives, etc. Briefly, the model contains four major compartments: *Body Blood*, *Thyroid*, *Liver*, and *Rest-of-Body* (*RB*) (Fig. 1A). The *Liver*, *RB*, and *Thyroid* compartments contain respective tissue blood (vasculature) and tissue proper (extra-vasculature) subcompartments. In each blood (sub)compartments, the binding events between THs and THBPs follow the law of mass action (Fig. 1B). As standard PBK practices, here the concentrations of molecular species in *Body Blood* are treated as their arterial concentrations (CA), and the concentrations in tissue blood are treated as proxies for their venous concentrations (CVT, CVRB, and CVL) (Fisher et al. 2020). The model produced several novel findings, including fast and near-equilibrium blood-tissue TH exchanges as an intrinsic robust mechanism against local metabolic perturbation and tissue influx as a limiting step for transient tissue uptake of THs in the presence of THBPs (Bagga et al. 2023). Since we were concerned with the steady-state behaviors of THs in local tissues, it was not necessarily to include the feedback regulation of TSH by THs in the model, which is a simplification that is also applied to the spatial model presented below.

**Figure 1.**
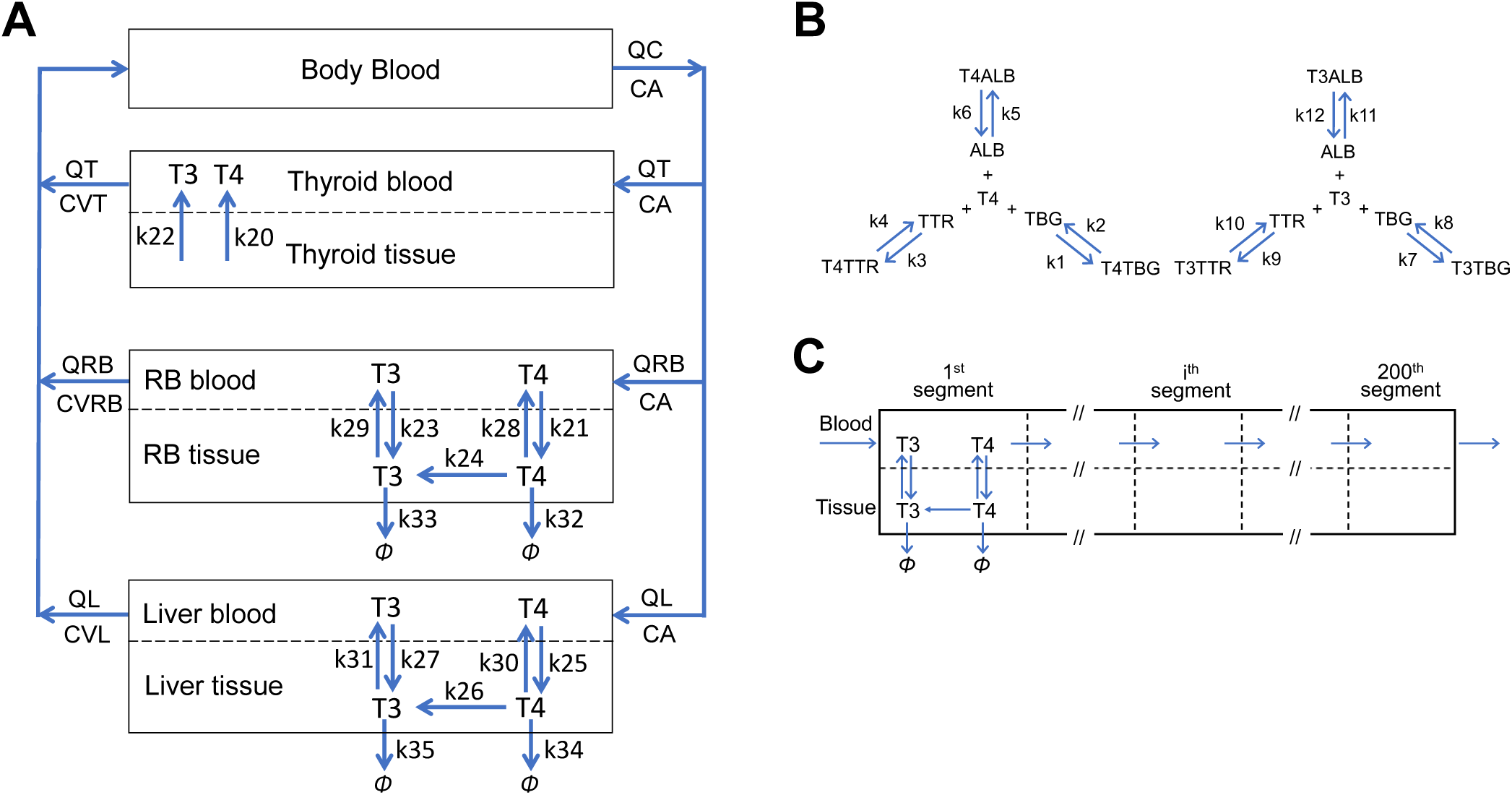
Schematic illustrations of the spatial human PBK model for T4 and T3. **(A)** The overall structure of the PBK model in (Bagga et al. 2023). *RB*: rest-of-body. QC: cardiac output (plasma portion); QT, QRB, and QL: rate of blood (plasma) flow to *Thyroid*, *RB*, and *Liver* respectively (QL combines blood flows from both the hepatic artery and portal vein); CA: arterial concentration of a TH species; CVT, CVRB, and CVL: venous concentration of a TH species in *Thyroid*, *RB*, and *Liver* respectively; *k*_20_ *– k*_35_: rate constants for production, transport, or metabolism processes as indicated. **(B)** Reversible binding of T4 and T3 with TBG, TTR and ALB. These binding events occur in the plasma of all blood (sub)compartments/subsegments. *k*_1_ *– k*_12_: rate constants for association and dissociation between THs and THBPs as indicated. Refer to (Bagga et al. 2023) for definitions and values of these parameters in (A) and (B). **(C)** Illustration of tissue segmentation in the spatial PBK model. The tissue influx, efflux, T4-to-T3 conversion, and metabolism occur in all segments, but are only illustrated in the 1^st^ segment.

### 2. Construction of the spatial human PBK model of THs

In order to examine the gradients of TH distribution in tissues, we extended the nonspatial PBK model above into a spatial model. Specifically, the *RB* and *Liver* compartments are divided into 200 linear segments of equal size with each segment containing its own tissue blood and tissue proper subsegments (Fig. 1C). 200 segments provide a sufficient resolution to simulate the concentration gradients as higher numbers of segments do not appreciably improve the precision (simulation results not shown). The concentration of each molecular specie in the last (200^th^) tissue blood subsegment is treated as its tissue venous blood concentration. Each subsegment is assumed to be well-mixed and has its own set of ordinary differential equations (ODEs) tracking the subsegment-specific rates of change of the variables. The method of lines is used to track the molecular species in each subsegment (Hamdi et al. 2007). The i^th^ and i+1^th^ blood subsegments are connected by the unidirectional blood flow of each molecular species. The kinetic constants *k*_1_ *– k*_12_ for TH and THBP binding in the blood (i.e., the association rate constants and dissociation rate constants) are assumed the same across all blood subsegments and equal to the values in the nonspatial PBK model. The bidirectional transport of free T4 and free T3 between the tissue blood and tissue proper subsegments as well as T4-to-T3 conversion metabolism and clearance metabolism within each tissue proper subsegment also occur as described in the nonspatial PBK model. The parameters describing the influx and efflux within each tissue segment (*k*_21_, *k*_23_, *k*_25_, *k*_27_, *k*_28_, *k*_29_, *k*_30_, and *k*_31_) are scaled from the values used in the nonspatial PBK model by dividing by 200, the total number of segments. The parameters describing the metabolism within each tissue segment (*k*_24_, *k*_26_, *k*_32_, *k*_33_, *k*_34_, and *k*_35_), which are first-order rate constants, remain the same as in the nonspatial PBK model. For homogenous simulations, the values of each of these parameters (*k*_21_, *k*_23_ – *k*_35_) are equal across all segments. For heterogeneous simulations where a parameter value may vary from the arterial to venous ends, a linear gradient was applied to each parameter across the segments such that the average is equal to the value used in the homogenous simulations.

One additional consideration is the possible bidirectional diffusion of molecular species across the blood subsegments and across the tissue subsegments. The diffusion of a molecular species would be normally modeled as (species_i-1_ + species_i+1_ -2 * species_i_)/dx^2^ where i represents the current subsegment, i-1 the subsegment before, i+ 1 the subsegment after, and *d*x is the length of the subsegment. Since the diffusion rate is much slower than the blood flow rate (Weisiger et al. 1986), including cross-subsegment diffusions did not produce any difference in the model’s behavior based on our simulations (results not shown). Therefore, for all spatial PBK simulations we proceeded without considering cross-subsegment diffusions.

### 3. Model parameters, equations, and simulation tools

Model parameter values and details of the source references and justifications, ordinary differential equations (ODEs), and algebraic equations were described in (Bagga et al. 2023). Both the nonspatial and spatial models were constructed in MATLAB R2019a (MathWorks Natick, Massachusetts, USA), and *ode15s* was used to numerically solve the ODEs. All MATLAB code will be available at https://github.com/pulsatility/2024-TH-PBK-Model.git.

## Results

### 1. Nonspatial PBK model

#### 1.1 TH unloading in *Liver blood*

Compared with the respective arterial concentrations, free T4 (*fT4*), *T4TBG*, *T4TTR*, and *T4ALB* in the hepatic venous blood leaving the *Liver* compartment are only negligibly lower, dropping by 0.02%-0.18% (Table S1). *fT4* drops by 0.18%, which is comparable to the percentage drop of *T4ALB* (0.17%), however, *T4TTR* drops only by 0.069% and *T4TBG* by 0.022%. This result suggests that only *T4ALB* is in near equilibrium with *fT4* as the blood transits through the *Liver*. Here, equilibrium refers to the state when the rate of association between *fT4* (or *fT3*) and either *TBG*, *TTR* or *ALB* equals the rate of dissociation of the respective TH-THBP complex. The difference in the percentage drops of these three T4-THBPs can be explained by the T4 residence time on these proteins relative to the blood transit time through the *Liver*. Given the vascular volume and blood flow rate of the *Liver*, as detailed in (Bagga et al. 2023), the blood transit time is about 8.3 seconds. The average residence time of T4 on TBG, TTR and ALB, as determined by the inverse of the dissociation rate constants *k*_2_, *k*_4_, and *k*_6_, is 55.55, 12, and 0.77 seconds respectively. Therefore, it is expected that *T4ALB* can unload quickly in response to the drop of *fT4* as *fT4* moves into the *Liver tissue*, while *T4TTR* unloads much more slowly, and *T4TBG* unloads the slowest during this short transit time. Despite these differences, the absolute amounts of T4 unloaded from the three THBPs are not much different – the concentration of *T4TBG* decreases the most, by 15.2 pM, followed by *T4ALB*, 12.3 pM, and *T4TTR*, 11.8 pM. This is because although the relative speed of T4 dissociation from TBG is much slower than from ALB, *T4TBG* has a much higher (nearly 10 times) abundance than *T4ALB*, resulting in a higher absolute amount of T4 unloaded in the *Liver blood* of the nonspatial model.

For T3 in *Liver*, the hepatic venous blood concentrations are negligibly lower than the arterial concentrations by 0.24%-0.42% (Table S1). Free T3 (*fT3*) drops by 0.42%, comparable to the percentage drop of *T3ALB* (0.4%), only slightly higher than *T3TTR* (0.36%), but tangibly higher than *T3TBG* (0.24%). The average residence time of T3 on TBG, TTR and ALB, as determined by the inverse of the dissociation rate constants *k*_8_, *k*_10_, and *k*_12_, is 6.06, 1.45, and 0.45 seconds respectively. Compared with the blood liver transit time of 8.3 seconds, it is clear that *T3ALB* and *T3TTR* can reach near equilibrium with *fT3* during this time, while *T3TBG* can only partially reach equilibrium. However, like the case of T4, *T3TBG* contributes the most T3 unloaded in the *Liver blood* with an absolute drop of 3.0 pM, followed by 1.3 pM by *T3ALB*, and a much smaller 0.3 pM by *T3TTR*.

#### 1.2 TH unloading in *RB blood*

For T4 in *RB*, the venous blood concentrations are negligibly lower than the arterial concentrations by 0.017%-0.078% (Table S1). Similar to the case in *Liver*, the percentage drop of *T4ALB* is comparable to that of *fT4*, while *T4TBG* drops by a much smaller fraction. Given the vascular volume and blood flow rate of *RB* (Bagga et al. 2023), the blood *RB* transit time is 20 seconds, which is much longer than the T4 residence time on ALB, nearly twice as long as on TTR, and less than half of the residence time on TBG. The absolute amounts of T4 unloaded from the three THBPs are quite different, where *T4TBG* unloads the most by 11.8 pM, *T4TTR* the second by 7.5 pM, and *T4ALB* the least by 5.6 pM.

For T3 in *RB*, the venous blood concentrations are negligibly lower than the arterial concentrations by 0.062%-0.088% (Table S1). *fT3* drops by 0.088%, similar to that of *T3ALB* and *T3TTR*, and only slightly higher than that of *T3TBG* (0.062%), which can be explained by the much longer blood *RB* transit time than the residence times of T3 on these THBPs (20 vs. 6.06, 1.45, and 0.45 seconds). Similar to T3 in *Liver*, *T3TBG* contributes nearly 70% of T3 unloaded in *RB blood* with an absolute decrease of 0.78 pM, followed by 0.29 pM by *T3ALB*, and a much negligible 0.07 pM by *T3TTR*.

### 2. Spatial PBK model

The nonspatial PBK model provides some preliminary insights into the relative contributions of THBPs to local TH delivery. A caveat of the nonspatial model is that the *Liver* and *RB* are treated as well-mixed compartments, so it is unable to predict the concentration gradients of THs in the tissues. To this end, we utilized the spatial PBK model, where both the *Liver* and *RB* compartments are simulated as 200 interconnected, consecutive segments as detailed in Methods (Fig. 1C). The half-lives of plasma T4 and T3 and the decay profiles in simulated T4 or T3 tracer experiments as previously described (Bagga et al. 2023), in either the presence or absence of THBPs, are nearly identical to the nonspatial model (simulation results not shown), indicating that the spatial model is equivalent to the nonspatial model as far as the overall TH kinetics are concerned.

#### 2.1 Overall contributions of THBPs to TH unloading in tissue blood

The spatial PBK model predicts that the arterial and venous plasma concentrations of both T4 and T3 are marginally lower than the corresponding concentrations in the nonspatial PBK model (Tables 1 vs. S1). For a given TH variable, the differences between the arterial and venous concentrations are similar between the two models, and the relative contributions of TBG, TTR and ALB to providing THs to the tissues are by and large in the same order as predicted by the nonspatial model. One exception is *Liver* T4, where although the contributions of the three THBPs are still comparable, ALB unloads (14.4 pM) 10% more than TBG (13.1 pM) does, while in the nonspatial model TBG unloads 20% more than ALB does. The percentage contributions by each THBP species in the spatial model are summarized in Fig. 2.

**Figure 2.**
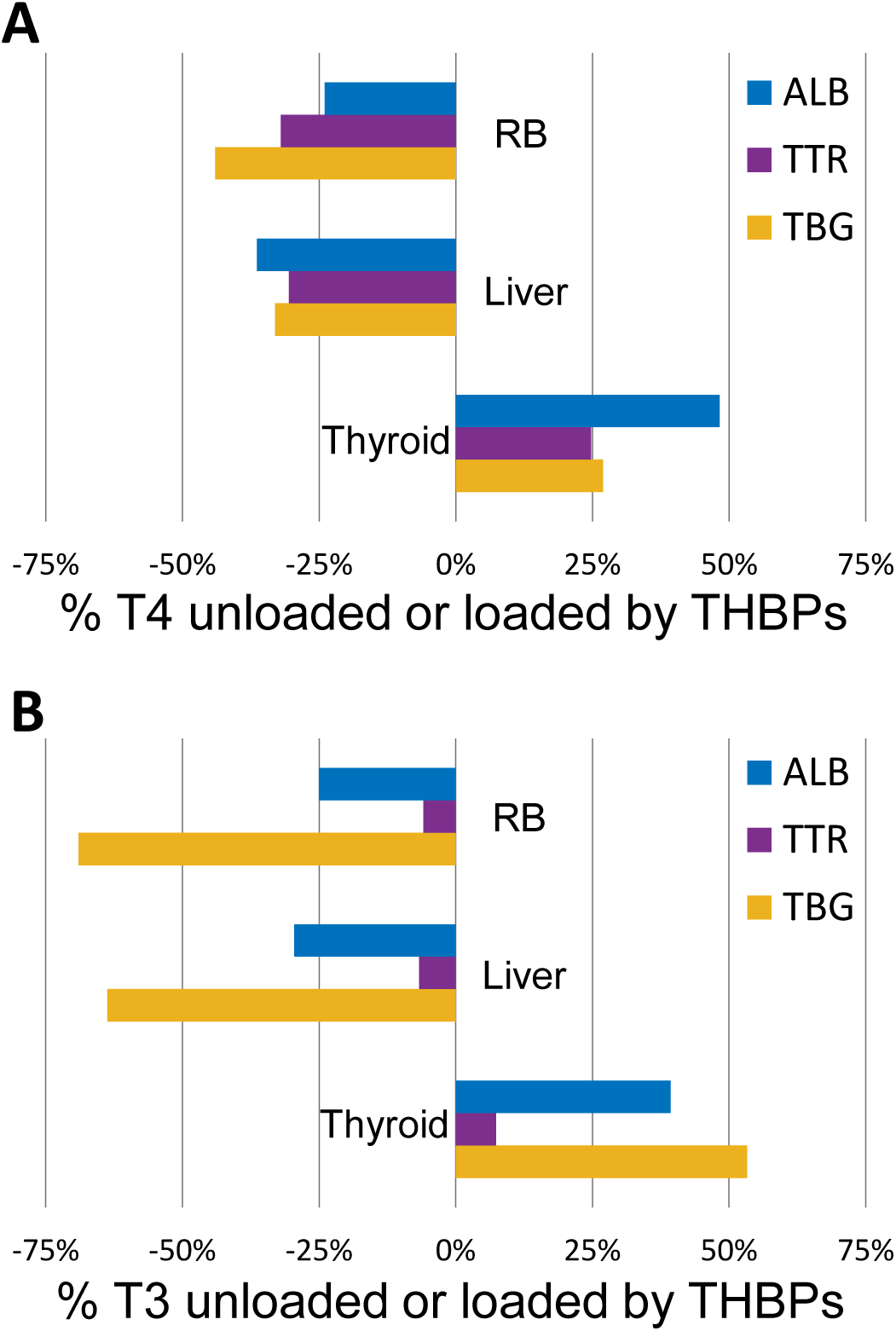
Percentage contributions of THBPs to loading and unloading of T4 (A) and T3 (B) in tissue blood in the spatial PBK model. Percentage contributions by ALB, TTR, and TBG in *RB*, *Liver*, and *Thyroid* compartments are indicated. Negative values denote TH unloading from THBPs and positive values denote TH loading onto THBPs.

#### 2.2 TH concentration gradients

The spatial plasma concentrations of *fT4* and the three T4-THBPs in tissue blood exhibit very different gradient profiles (Fig. 3A-3E). *fT4* (Fig. 3A) and *T4ALB* (Fig. 3D) in both *Liver blood* and *RB blood* show similar, exponential-like drops (with the exception that *fT4* exhibits a discrete drop in the first segment). When examined on logarithmic scale of the Y axis, the decays do not follow a straight line (results not shown), indicating that they are not truly exponential. In contrast, *T4TBG* exhibits a decreasing trend that is concave downward (Fig. 3B), while *T4TTR* is concave downward initially but then transitions to a nearly linear decay phase, especially in *RB blood* (Fig. 3C). Interestingly, *Total T4* decays in a basically linear fashion (Fig. 3E). The concentration gradients of T4 in *Liver tissue* and *RB tissue* follow the same trend and profile as *fT4* in the respective tissue blood, and the percentage drops from the first segment to the last segment, which are negligibly small, are also the same as the drop of *fT4* (results not shown). These gradient profiles are qualitatively similar in both the *Liver* and *RB* compartments, but the concentration drops are generally steeper in *Liver* than in *RB*. The gradient profiles of T3 are similar to those of T4 (Fig. 3F-3J), except that *T3TTR* exhibits a concave-upward profile for the most part (Fig. 3H) whereas *T4TTR* is more concave-downward (Fig. 3C).

**Figure 3.**
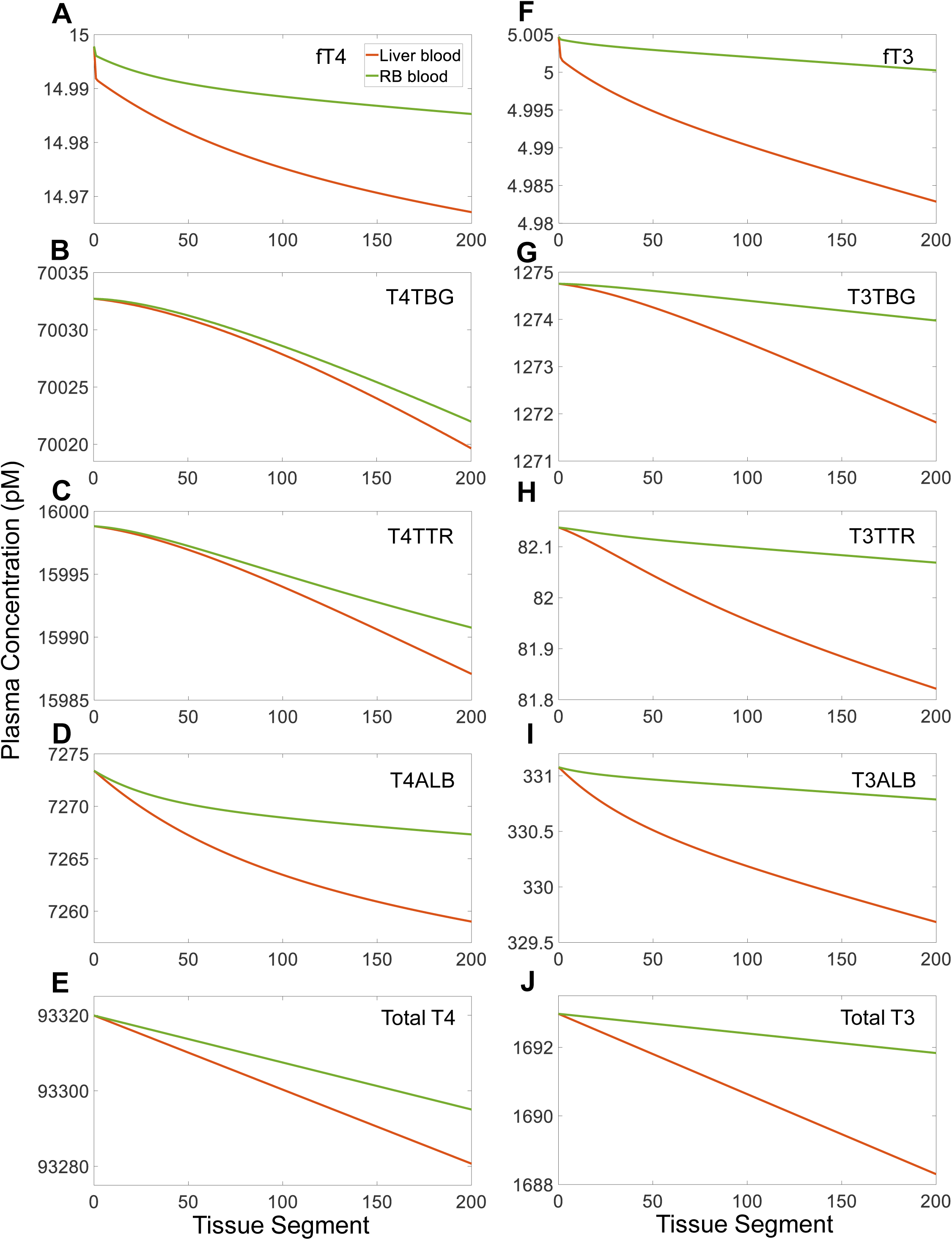
Concentration gradients of free THs, TH-THBPs, and total THs in tissue blood in the spatial PBK model. **(A-E)** Plasma concentrations of *fT4*, *T4TBG*, *T4TTR*, *T4ALB*, and *Total T4* in *Liver blood* (orange) and *RB blood* (green) respectively. **(F-J)** Plasma concentrations of *fT3*, *T3TBG*, *T3TTR*, *T3ALB*, and *Total T3* in *Liver blood* (orange) and *RB blood* (green) respectively. Concentrations in segment 0 represent the arterial concentrations and the concentrations in segment 200 represent the venous concentrations.

#### 2.3 Nonlinear, spatially dependent THBP unloading of THs in tissue blood

We next calculated the difference in T4-THBP concentrations between two consecutive tissue blood segments, which represents the amount of T4 that is unloaded by the THBPs in the downstream segment of the two. It is interesting to note that the concentration drop of each THBP species varies depending on the location of the segments (Fig. 4A and 4B). ALB clearly contributes the most T4 in the upstream *Liver* segments, but as the blood perfuses downstream, the contributions by TBG and TTR start to catch up halfway through. Toward the venous end of the *Liver* compartment, TBG becomes the dominant contributor, TTR the intermediate contributor, and ALB the smallest contributor (Fig. 4A). A similar change of order in contribution between ALB, TBG and TTR also occurs in the *RB* compartment, although more upstream, nearly quarter-way through (Fig. 4B). Interestingly, the *T4TTR* drop exhibits a nonmonotonic profile in *RB*. The contributions of TBG and ALB to T3 unloaded in *RB blood* exhibit similar profiles to the case of T4, with changes in the order of contribution occurring at further upstream locations (Fig. 4C and 4D). *T3TTR* has the smallest contribution also showing nonmonotonicity in the *Liver blood* and *RB blood* compartments. At downstream locations in *RB*, the amount of T3 unloaded from each of the three THBPs seems to reach a constant.

**Figure 4.**
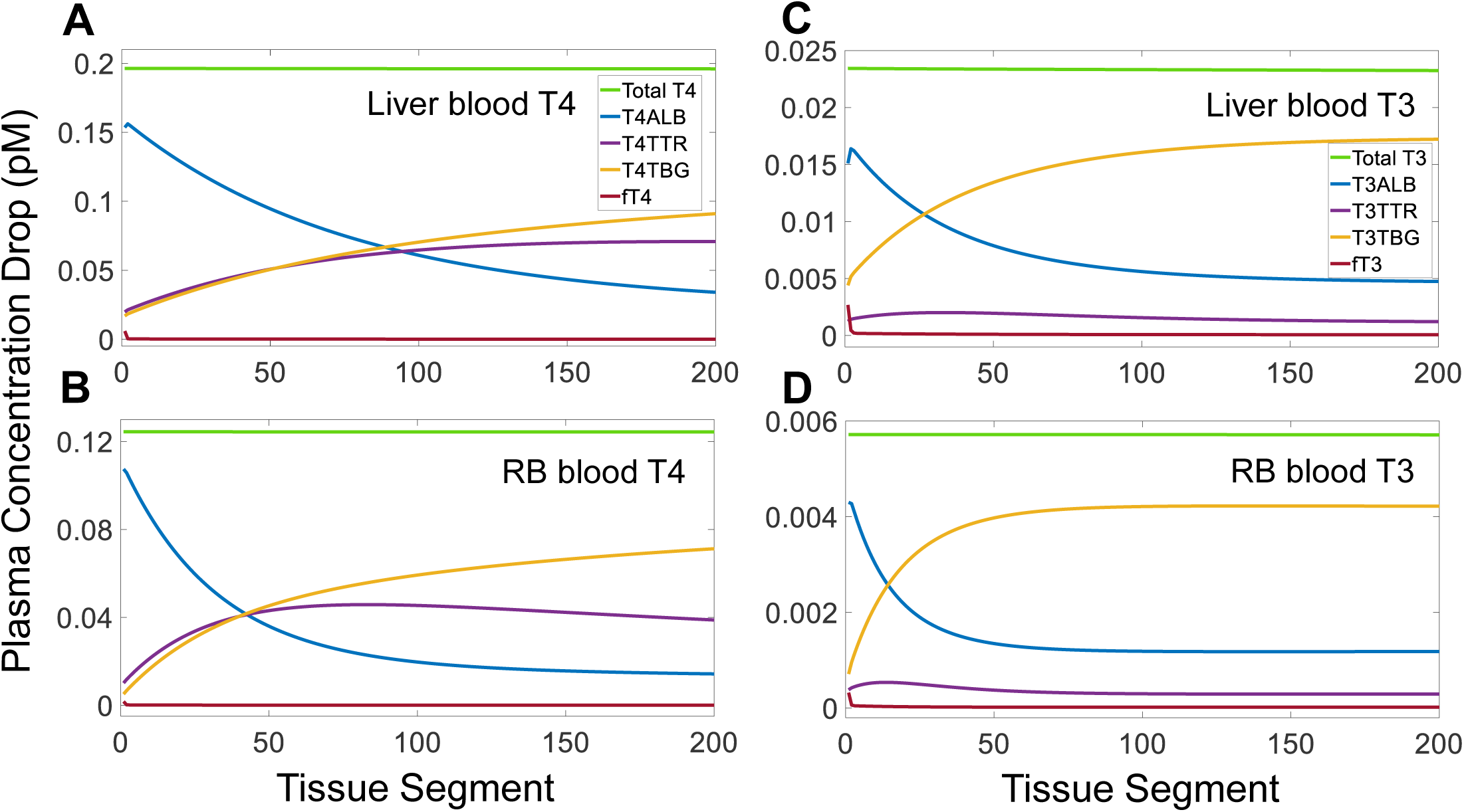
Segment-to-segment net differences in concentrations of free THs, TH-THBPs, and total THs in tissue blood in the spatial PBK model. The value of concentration drop for each segment is calculated by subtracting the concentration in the current segment from that in the immediate upstream segment. For the drop in the first segment, the arterial blood concentration (segment 0) is used as the upstream concentration. **(A-B)** Segment-to-segment drops in plasma concentrations of *fT4*, *T4TBG*, *T4TTR*, *T4ALB*, and *Total T4*, as indicated in (A), in *Liver blood* and *RB blood* respectively. **(C-D)** Segment-to-segment drops in plasma concentrations of *fT3*, *T3TBG*, *T3TTR*, *T3ALB*, and *Total T3*, as indicated in (C), in *Liver blood* and *RB blood* respectively.

This nonlinear, spatially dependent, differential TH unloading by the three THBPs is due, at least in part, to the differences in the residence time of T4 and T3 on the THBP molecules. In the upstream segments, for instance, as *fT4* moves into the tissue such that its local concentration drops, *T4ALB* releases T4 much more quickly than *T4TTR* and *T4TBG*. The T4 released by *T4ALB* likely further slows down the release of T4 from *T4TTR* and *T4TBG*, resulting in the concave-downward gradients of the two species (Fig. 3B and 3C). As the blood flows through the remaining segments downstream where *fT4* drops to lower levels, *T4TTR* and *T4TBG* are further removed from their equilibrium with *fT4* (Fig. 5A-5B and 5D-5E). This built-up “tension” or disequilibrium begins to force *T4TTR* and *T4TBG* to release more T4 than in the upstream segments. While in contrast, *T4ALB* moves further toward equilibrium, therefore releasing less T4 (Fig. 5C and 5F). The evolution of the binding disequilibrium between T3 and THBPs through tissue blood shows similar trends to the case of T4 (Fig. 6). *T3TTR* binding disequilibrium exhibits nonmonotonic changes in both *Liver blood* (Fig. 6B) and *RB blood* (Fig. 6E), similar to *T4TTR* in *RB blood* (Fig. 5E), which explains the bell-shaped contributions to T3 (Fig. 4C-4D) and T4 (Fig. 4B) by TTR.

**Figure 5.**
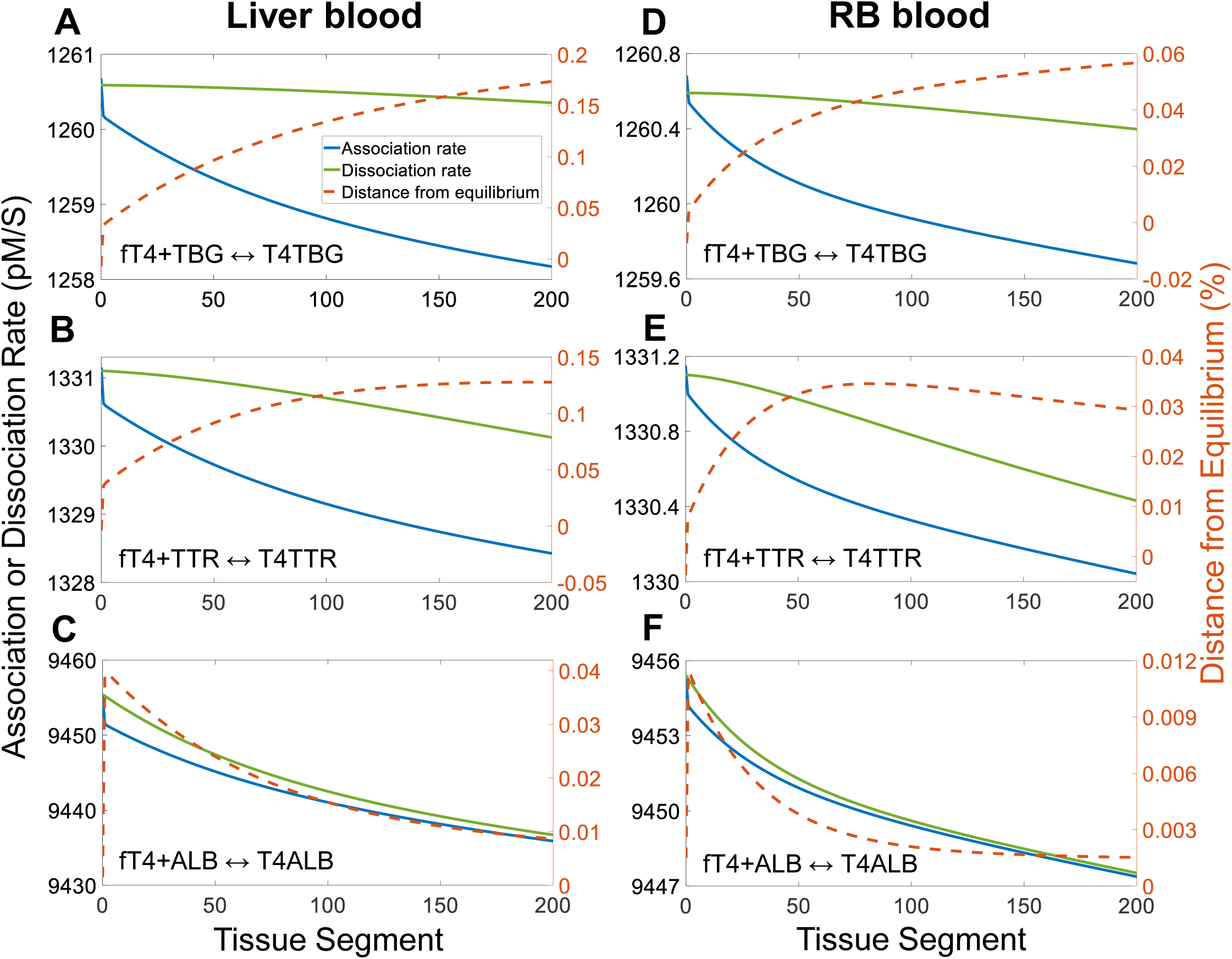
Evolution of the binding kinetics between T4 and THBPs through tissue blood in the spatial PBK model. *Association rate* (solid blue), *Dissociation rate* (solid green), and *Distance from equilibrium* (dashed orange) for *fT4* binding to *TBG* **(A)**, *TTR* **(B)**, and *ALB* **(C)** in *Liver blood*. *Association rate*, *Dissociation rate*, and *Distance from equilibrium* for *fT4* binding to *TBG* **(D)**, *TTR* **(E)**, and *ALB* **(F)** in *RB blood*. *Distance from equilibrium* = (*Dissociation rate* - *Association rate*) ⁄ *Association rate* × 100%. Segment 0 represents the arterial blood. The same color scheme is used for all panels as indicated in (A).

**Figure 6.**
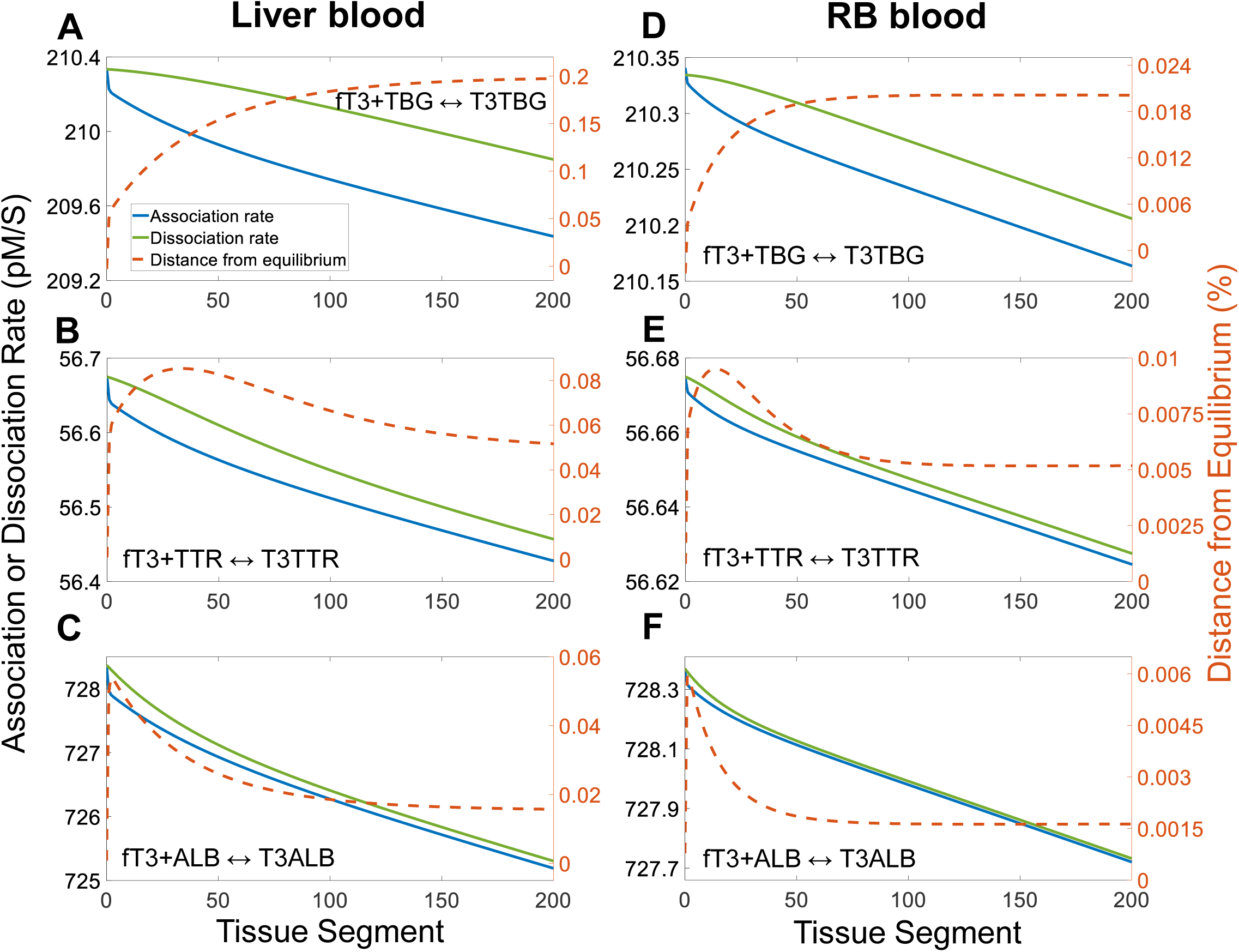
Evolution of the binding kinetics between T3 and THBPs through tissue blood in the spatial PBK model. *Association rate* (solid blue), *Dissociation rate* (solid green), and *Distance from equilibrium* (dashed orange) for *fT3* binding to *TBG* **(A)**, *TTR* **(B)**, and *ALB* **(C)** in *Liver blood*. *Association rate*, *Dissociation rate*, and *Distance from equilibrium* for *fT3* binding to *TBG* **(D)**, *TTR* **(E)**, and *ALB* **(F)** in *RB blood*. *Distance from equilibrium* = (*Dissociation rate* - *Association rate*) ⁄ *Association rate* × 100%. Segment 0 represents the arterial blood. The same color scheme is used for all panels as indicated in (A).

The differential contribution to the amounts of THs unloaded in tissue blood by different THBPs cannot be explained, however, solely by the difference in the residence time of T4 or T3 on THBP molecules. There is a quantitative mismatch. For instance, the residence time of T4 on ALB is 70 times shorter than on TBG and nearly 16 times shorter than on TTR. However, in the first segment of the *Liver*, *T4ALB* contributes only about 7 times higher T4 than *T4TBG*, and the *T4TBG* contribution is on par with *T4TTR* contribution (Fig. 4A). An examination of the absolute association rates and dissociation rates of all TH/THBP pairs reveals that the relative contributions to THs unloaded by the three THBPs in the most upstream segments are basically proportional to the absolute dissociation rates in the arterial blood (Figs. 5 and 6 and Table S2). These dissociation rates are determined not only by the dissociation rate constant parameters (which determine the residence time), but also by the concentrations of TH-THBP complexes. In the case of contributions to the amount of T4 unloaded in local tissues, *T4TBG* and *T4TTR* have higher concentrations than *T4ALB*, which compensate to a great extent their longer residence time.

Despite the nonlinear, varying contribution by each of the three THBPs across tissue segments, the total amounts of T4 and T3 unloaded by all three THBPs combined is highly constant across tissue segments (Fig. 4, green lines). The amount of T4 unloaded and thus delivered in the last segment is only 0.16% and 0.07% lower than that in the 1^st^ segment in the *Liver* and *RB* compartments respectively, and in the case of T3, the difference is 0.82% and 0.12% respectively. These results support the notion that THBPs play a crucial role in maintaining a constant concentration and uniform delivery of THs through the tissues. This is further demonstrated when all three THBPs are removed in the spatial model (Fig. 7). The arterial concentration of *fT4* increases dramatically to 34 pM, compared with 15 pM when all three THBPs are present, and the venous concentrations leaving the *Liver* and *RB* compartments drop to 2.5 and 6.5 pM respectively, forming steep exponential gradients (Fig. 7A and 7B, solid line). T4 concentrations in the *Liver tissue* and *RB tissue* follow a similar descending gradient with nearly identical percentage drops as *fT4* (Fig. 7C and 7D, solid line).

**Figure 7.**
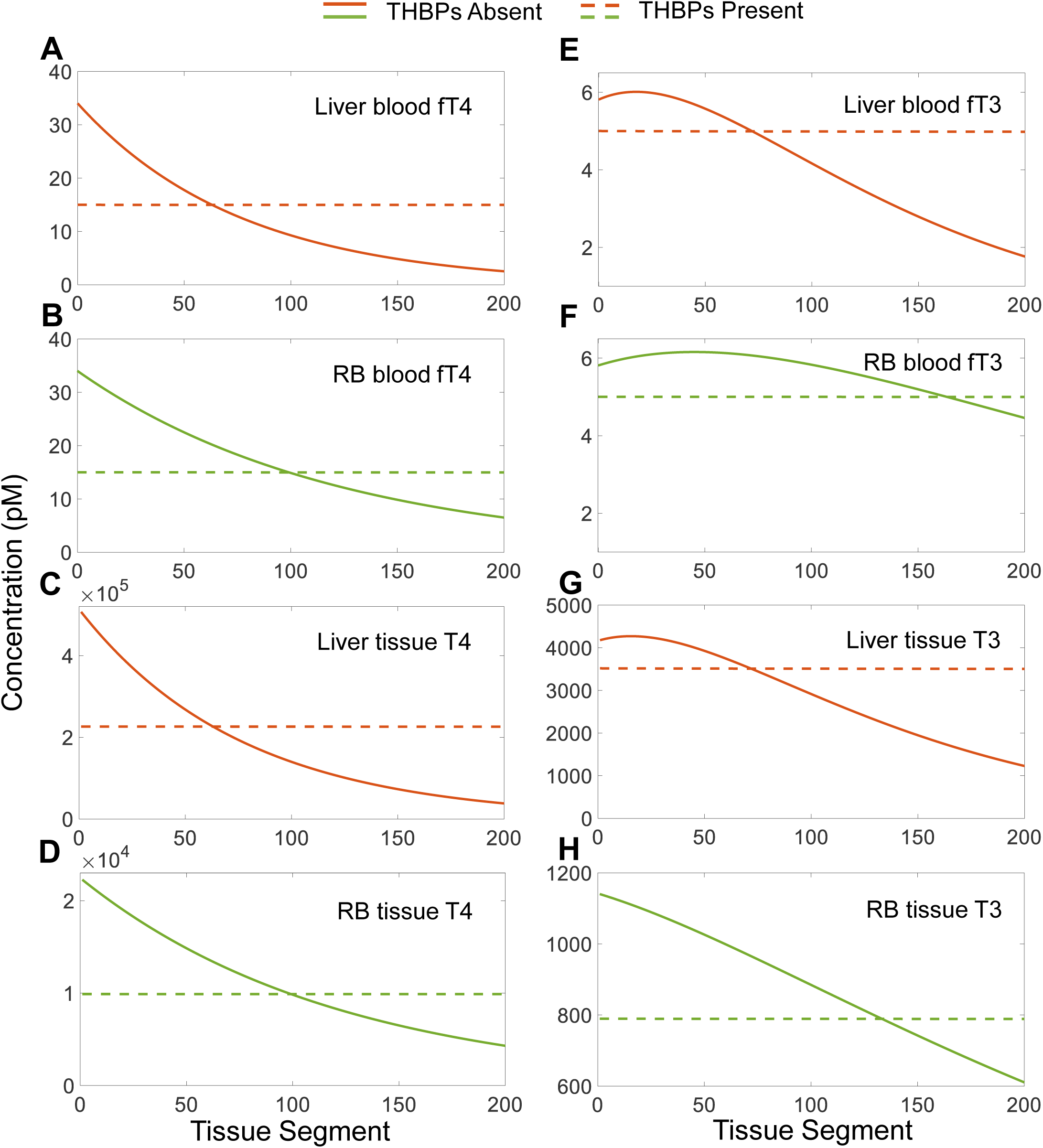
Concentration gradients of THs in blood and tissues in the absence all THBPs in the spatial PBK model. **(A-B)** Plasma concentrations of *fT4* in *Liver blood* and *RB blood* respectively. **(C-D)** Concentrations of T4 in *Liver tissue* and *RB tissue* respectively. **(E-F)** Concentrations of *fT3* in *Liver blood* and *RB blood* respectively. **(G-H)** Concentrations of T3 in *Liver tissue* and *RB tissue* respectively. Solid line: all THBPs absent, dashed line: all THBPs present. Concentrations in segment 0 represent the plasma concentrations in arterial blood.

The arterial concentration of *fT3* increases to 5.81 pM, compared with 5 pM when all three THBPs are present. Surprisingly, unlike T4 which exhibits a monotonically descending gradient, plasma *fT3* exhibits a bell-shaped gradient in both *Liver* and *RB* when all three THBPs are absent (Fig. 7E and 7F, solid line). *fT3* first increases slightly less than 1/10^th^ way through the *Liver blood*, and then decreases in a nearly linear fashion followed by a slightly concave-upward decay down to 1.76 pM (Fig. 7E). The bell shape of the plasma *fT3* gradient in *RB* is more dramatic, where *fT3* first rises to 6.16 pM more than 1/5^th^ way through the tissue, and then decreases to 4.46 pM as it exits the tissue (Fig. 7F). T3 in *Liver tissue* follows a similar nonmonotonic gradient profile as *fT3* in *Liver blood* (Fig. 7G, solid line) but T3 in *RB tissue* only exhibits a monotonic decay (Fig. 7H, solid line). The nonmonotonicity of *fT3* gradients in the tissue blood in the absence of all three THBPs is explained in Fig. S1.

The above results clearly demonstrate that THBPs are essential to maintain constant concentrations and uniform delivery of THs through the tissues. We next examined if any one of the three THBPs is sufficient to fulfill this function. When only either TBG, TTR, or ALB remains in the model, the concentrations of T4 and T3 across the blood and tissue segments exhibit negligible drops and the arterial concentrations are nearly identical to the concentrations when all three THBPs are present (Figs. S2-S4). The only notable changes are *fT3* concentration when only TTR is present, where it drops from 5.2 pM in the arterial blood to 4.9 pM in *Liver* venous blood (Fig. S3C). This is because *T3TTR* is the least abundant among the three T3-THBPs, thus a large percentage drop is needed to meet the demand by *Liver tissue*. Interestingly, just like the concentration gradients of *Total T4* and *Total T3* when all three THBPs are present, the concentration gradients of *T4TBG* and *T3TBG*, *T4TTR* and *T3TTR*, or *T4ALB* and *T3ALB* (which are essentially the total T4 and T3 respectively since only a single THBP species is present in each case) are also basically linear and exhibit the same absolute drop (e.g., about 40 pM for T4 and 4.5 pM for T3 from segment 0 to segment 200 in *Liver blood*). These results indicate that any of the three THBPs is sufficient to ensure uniform delivery of THs in tissues.

Lastly, we examined the situation when only one THBP is absent. With a combination of any two THBPs present, the concentration gradients of *Total T4* and *Total T3* are also basically linear and exhibit the same absolute drops as when all three THBPs are present, indicative of uniform delivery of THs in tissues (Figs. S5-S7). *fT4* and the T4-THBP species that has the shorter residence time for T4 always exhibit concave-upward decay profiles, while the T4-THBP species that has the longer residence time exhibits a concave-downward decay profile. The combination of ALB and TBG produces the most contrasted profiles (Fig. S6).

In our model, RB is a lumped compartment of all extrahepatic and extrathyroidal tissues, thus its behavior represents the average of these tissues. However, an individual tissue can be either more rapidly or more slowly perfused than the average. Moreover, the local production and metabolism of T3 can vary depending on the relative expression/activity levels of DIO2 and DIO3 such that the tissue can be either a T3 source or T3 sink (Goemann et al. 2018). We next explored how these different local conditions can affect THBP unloading or loading of THs. For simplicity, this was done by varying relevant parameters in the *RB* compartment while clamping all state variables in *Body Blood* at their respective baseline levels so that the local parameter changes do not alter the systemic TH levels. From the perspective of examining the roles of THBPs in TH unloading and loading, this is equivalent to splitting a small tissue compartment from *RB* and varying relevant parameters. When the *RB* blood flow rate is increased by 4 fold and there is no local T4-to-T3 conversion (*k*_24_=0) and thus the tissue acts as a pure T3 sink, all T4 (Fig. 8A-8E, solid red line) and T3 (Fig. 8F-8J, solid red line) species in *RB blood* drop less than at the default (average) condition. ALB unloads the majority of T4 while TBG and TTR make comparable contributions (Fig. 9A); the contribution to T3 unloading follows the order of T3-THBP abundances, i.e., T3TBG>T3ALB>T3TTR (Fig. 9B). Conversely, when the blood flow rate is decreased by 4-fold and local T4-to-T3 conversion still absent, all T4 and T3 species (dashed line) drop more than in the default condition. The contribution to TH unloading follows the order of TH-THBP abundances: T4TBG>T4TTR>T4ALB for T4 (Fig. 9C) and T3TBG>T3ALB>T3TTR for T3 (Fig. 9D). When the blood flow rate is increased by 4-fold and there is no local T3 degradation (*k*_33_=0) and thus the tissue acts as a pure T3 source, the T4 species have the same concentration profiles (Fig. 8A-8E, solid light blue line) as when the blood flow rate is also 4 times faster but *k*_24_=0 (solid red). In contrast, all T3 species exhibit somewhat increasing concentration gradients (Fig. 8F-8J, solid light blue line), indicating that THBPs load as opposed to unload T3 under this condition. This is reflected by the negative segment-to-segment concentration drops (Fig. 9F), and the contribution to T3 loading follows the order of T3-THBP abundances, i.e., T3TBG>T3ALB>T3TTR. Lastly, when the blood flow rate is decreased by 4-fold and local T3 degradation still absent, there is no effect on T4 (Fig. 8A-8E, dashed light blue line), similar to when the blood flow rate is also 4 times slower but *k*_24_=0 (dashed red). However, all T3 species increase dramatically as they flow through the tissue (Fig. 8F-8J, dashed light blue line), and the contribution to T3 loading also follows the order of T3-THBP abundances (Fig. 9H).

**Figure 8.**
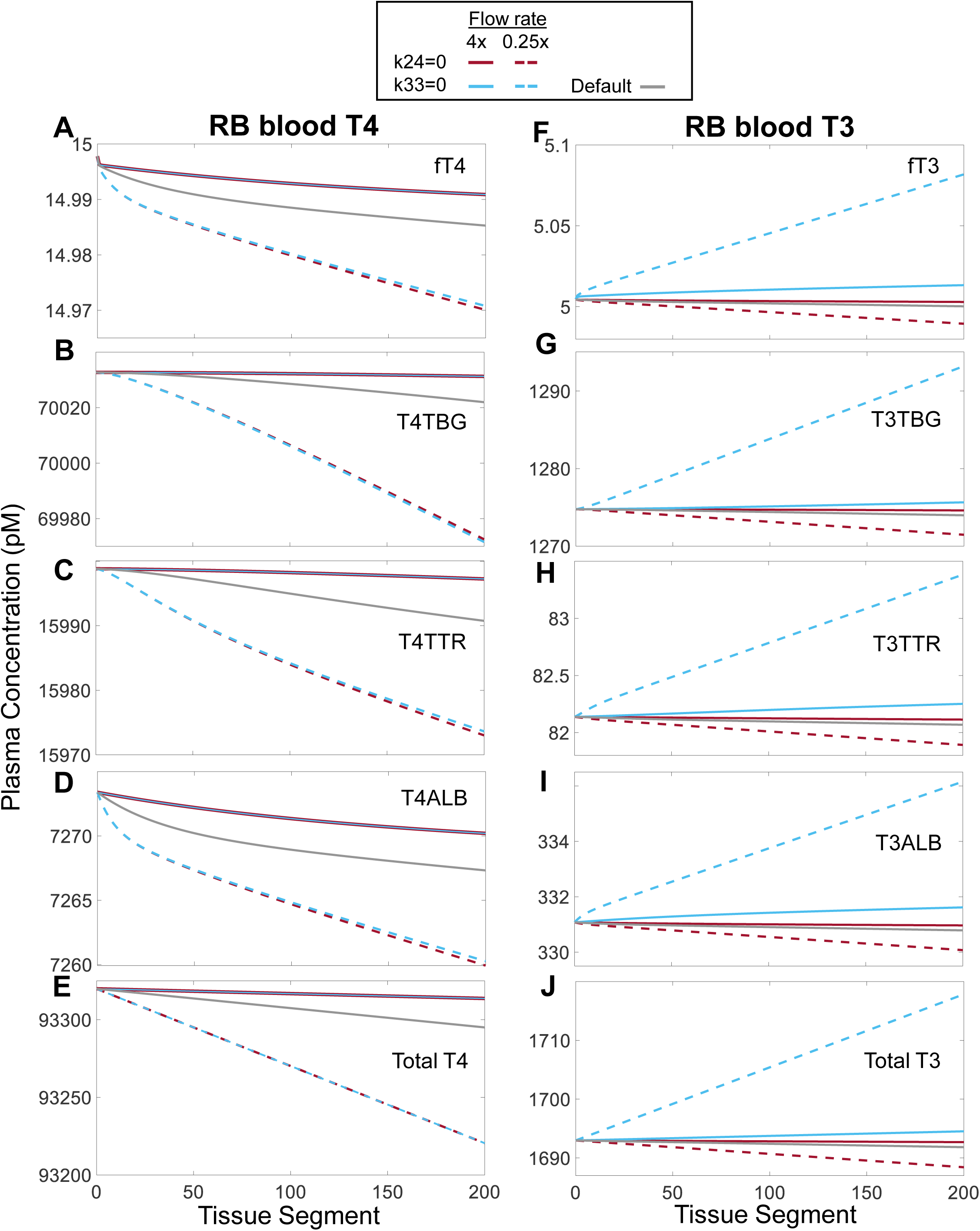
Concentration gradients of various TH species in *RB blood* in the spatial PBK model at different blood flow rates and *RB tissue* T3-related parameter values. (A-E) Plasma concentrations of *fT4*, *T4TBG*, *T4TTR*, *T4ALB*, and *Total T4* in *RB blood*. **(F-J)** Plasma concentrations of *fT3*, *T3TBG*, *T3TTR*, *T3ALB*, and *Total T3* in *RB blood*. The simulations were done by first clamping the plasma concentrations of all variables in *Body Blood* at basal steady-state levels as in Table 1, and then either increasing by 4-fold (4x) or decreasing by 4-fold (0.25x) the blood flow rate (QRB) from default value and setting either *k*_24_=0 (*k*_32_ was increased accordingly to keep overall *RB tissue T4* metabolism unchanged) or *k*_33_=0 as indicated on the top.

**Figure 9.**
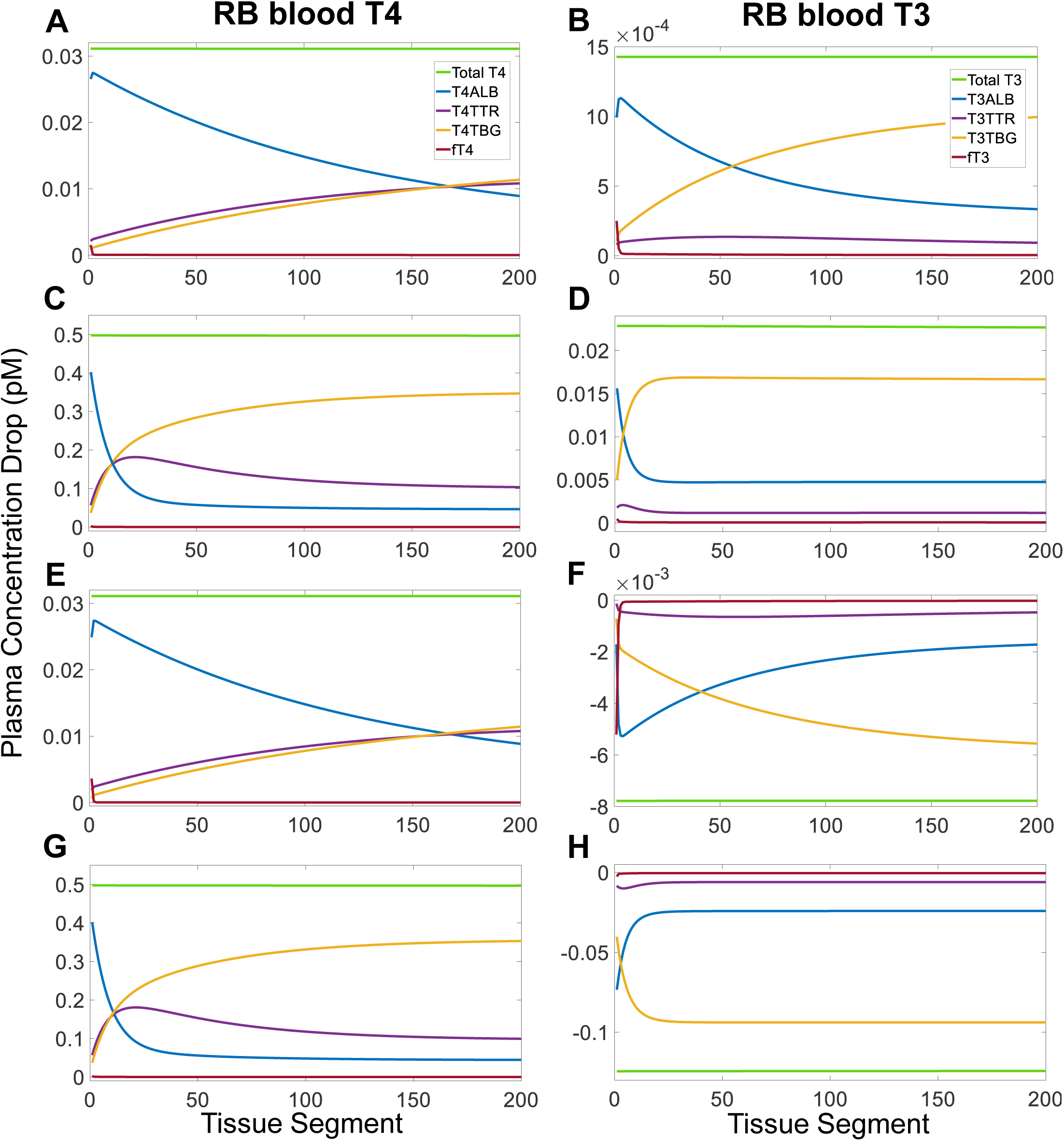
Segment-to-segment net differences in concentrations of free THs, TH-THBPs, and total THs in *RB blood* in the spatial PBK model at different blood flow rates and *RB tissue* T3-related parameter values. The value of the concentration drop for each segment is calculated as detailed in Fig. 4 legend, and the simulations were conducted as detailed in Fig. 8 legend. **(A-B)** QRB=4xdefault and *k*_24_=0, **(C-D)** QRB=0.25xdefault and *k*_24_=0, **(E-F)** QRB=4xdefault and *k*_33_=0, and **(G-H)** QRB=0.25xdefault and *k*_33_=0.

**Table 1.**
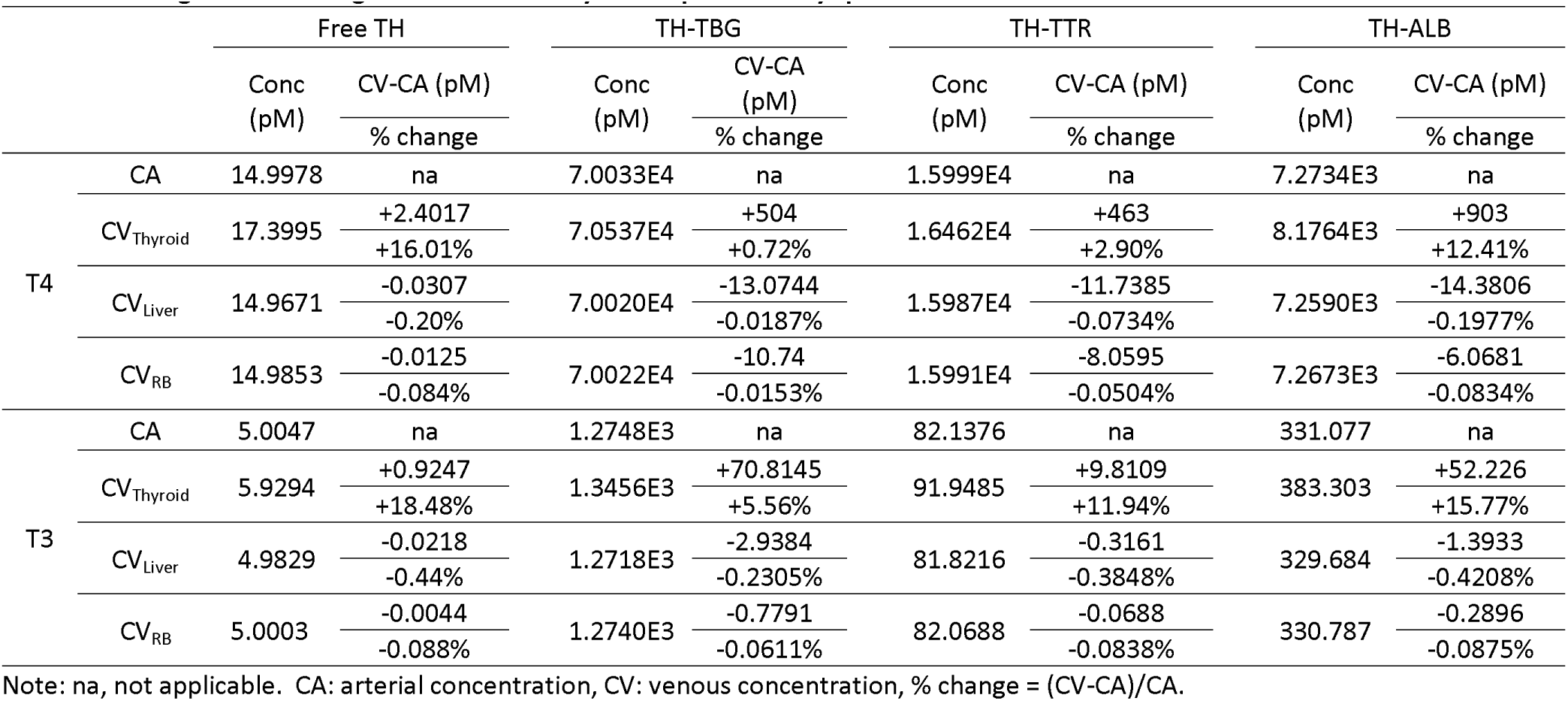
Loading and unloading of THs in tissues by THBPs predicted by spatial PBK model.

#### 2.4 Heterogeneous transport and metabolism across tissue segments

The above spatial analysis has the underlying assumption that each of the cross-membrane transport and metabolism parameters remains constant across the tissue segments from the arterial to venous ends. However, this assumption may not be true for all tissues or organs. For instance, the liver is known for its zonal heterogeneity in gene expression (Gebhardt and Matz-Soja 2014). Thus, transporters such as MCT10 as well as TH-metabolizing enzymes such as DIO1, UGT and SULT might have possible gradients from the periportal to central vein layers within the liver lobule (Halpern et al. 2017). In this section, we explored hypothetically how such intra-tissue heterogeneity alters THBP contribution to TH tissue delivery and TH tissue concentrations. For simplicity, a 10-fold linear gradient (either increasing or decreasing) was applied to one parameter (*k*_21_, *k*_23_ – *k*_35_) at a time with the average equal to the default value used in the homogenous case above. Below we reported the results for a select few parameters for the *Liver* and the results for the remaining parameters are provided in Supplemental Material.

When *k*_25_, the rate constant for T4 liver influx, increases from segment 1 through 200 (Fig. 10A), *fT4* in *Liver blood* barely drops percentage-wise, exhibiting a small concave-down gradient (Fig. 10B, red line) as compared with the concave-up gradient when *k*_25_ is constant (Fig. 10B, gray line). *Total T4* in *Liver blood* decreases with a concave-down gradient vs. a linear gradient with constant *k*_25_, but by the same amount in both cases (Fig. 10C). As far as T4 contributions by THBPs are concerned, they are all spatially dependent, but their nonlinearities are modulated by *k*_25_ variation as compared with constant *k*_25_ (Fig. 10E vs. Fig. 4A). ALB remains as the highest contributor throughout most segments, while TBG and TTR have similar but lower contributions. The total amount of T4 unloaded by all three THBPs combined increases linearly across the segments. Correspondingly, T4 in *Liver tissue* increases linearly as well, by nearly 10-fold (Fig. 10D). The kinetic changes of T4 caused by the *k*_25_ gradient also affect T3. *fT3* in *Liver blood* exhibits a small nonmonotonic profile (Fig. 10G) and *Total T3* also has a U-shaped profile vs. a linear one with constant *k*_25_, but it drops by the same amount in both cases (Fig. 10H). TBG still contributes to T3 the highest throughout most segments particularly downstream ones followed by ALB, and interestingly ALB reverts to net loading of T3 toward the venous end (Fig. 10J). The total amount of T3 unloaded by all three THBPs combined decreases linearly across the segments with net loading toward the venous end. Despite that T4 in *Liver tissue* increases by 10-fold across the segments, T3 in *Liver tissue* is insensitive to the change, increasing only by 3% (Fig. 10I). A decreasing *k*_25_ gradient has opposite effects compared with an increasing *k*_25_ gradient. When *k*_30_, the rate constant for T4 liver efflux, varies from segment 1 through 200, it has an opposite effect to varying *k*_25_ (Fig. 11). It is worth noting that despite linear *k*_30_ variation, T4 and T3 in *Liver tissue* exhibit highly hyperbolic-like gradients (Fig. 11D and 11I). As with varying *k*_25_, Liver tissue T4 is sensitive to *k*_30_ changes but T3 is much less sensitive. The nonlinear T4 and T3 gradients are explained as follows. Because the influx and efflux operate near equilibrium and the *fT4* gradient from the arterial to venous ends is minimal (Fig. 11B), the influx rate remains largely constant across segments. Therefore, when *k*_30_ varies linearly, for the efflux rate (which is equal to *k*_30_\**T4LT*\**fu_T4LT_*) to match the constant influx rate, mathematically, T4 in *Liver tissue* (*T4LT*) has to change inversely, resulting in a hyperbolic-like shape. This nonlinearity results in unequal metabolism of T4 in the first 100 segments vs. the last 100 segments, such that the overall T4 metabolism in the *Liver tissue* is higher than when *k*_30_ is constant, resulting in a greater drop in *Total T4* levels (Fig. 11C). Because of the T4-to-T3 conversion, T3 in *Liver tissue* has a similar nonlinear profile to T4 in *Liver tissue* with a much smaller gradient (about 9%).

**Figure 10.**
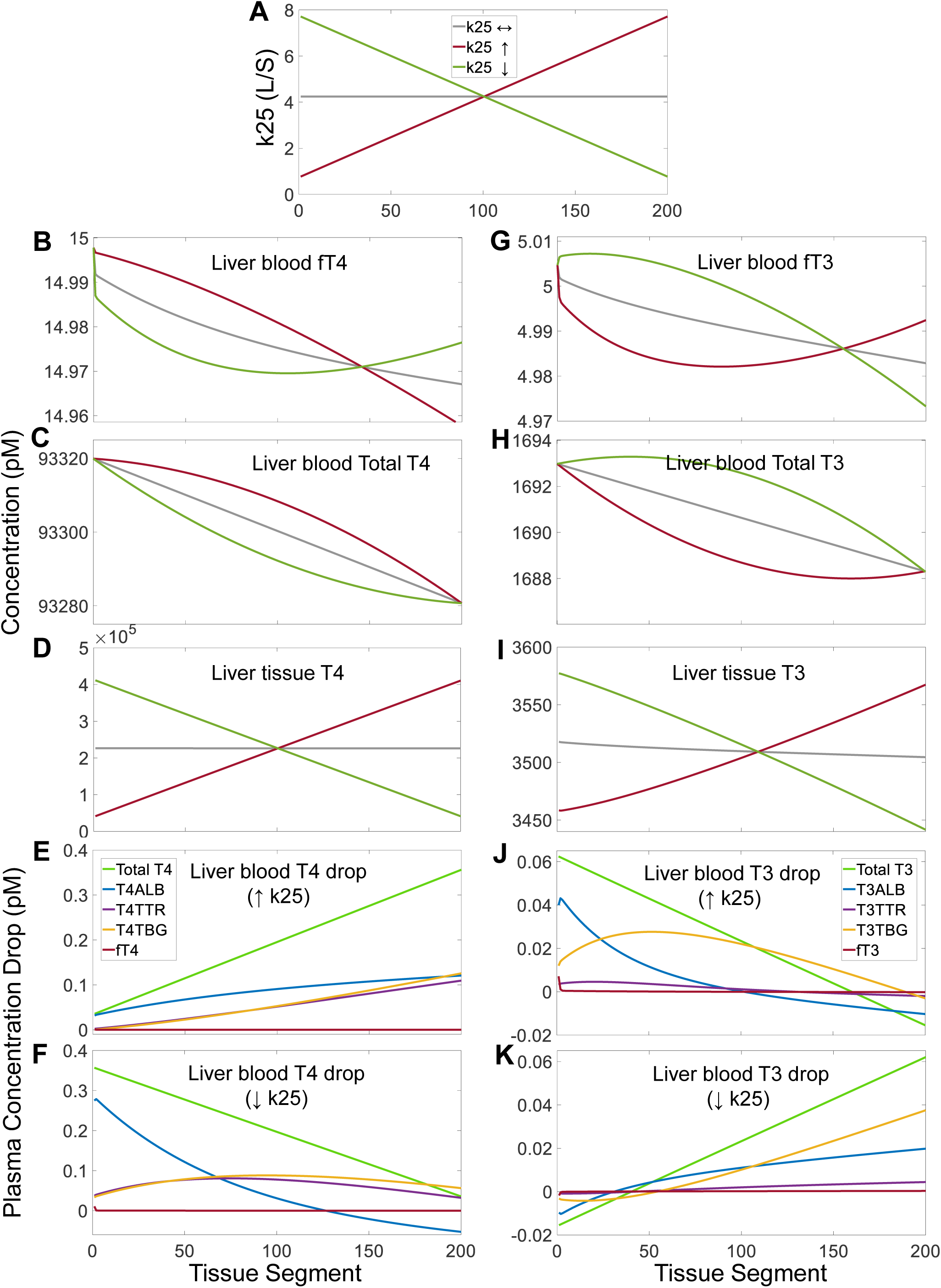
Effects of *k*_25_ gradients across *Liver* segments on TH concentrations in *Liver* and contributions of THBPs to TH loading and unloading in the spatial PBK model. (A) Linearly increasing, decreasing, or constant *k*_25_ gradients implemented as indicated. **(B-D)** T4 concentrations in *Liver blood* and *Liver tissue* as indicated. **(E-F)** Segment-to-segment net differences in plasma concentrations of *fT4*, *T4TBG*, *T4TTR*, *T4ALB*, and *Total T4* as indicated in *Liver blood* for increasing and decreasing *k*_25_ gradients respectively. **(G-I)** T3 concentrations in *Liver blood* and *Liver tissue* as indicated. **(J-K)** Segment-to-segment net differences in plasma concentrations of *fT3*, *T3TBG*, *T3TTR*, *T3ALB*, and *Total T3* as indicated in *Liver blood* for increasing and decreasing *k*_25_ gradients respectively. The same color scheme is used for panels (B-D, G-I) as indicated in (A), for panel (F) as in (E), and for panel (K) as in (J).

**Figure 11.**
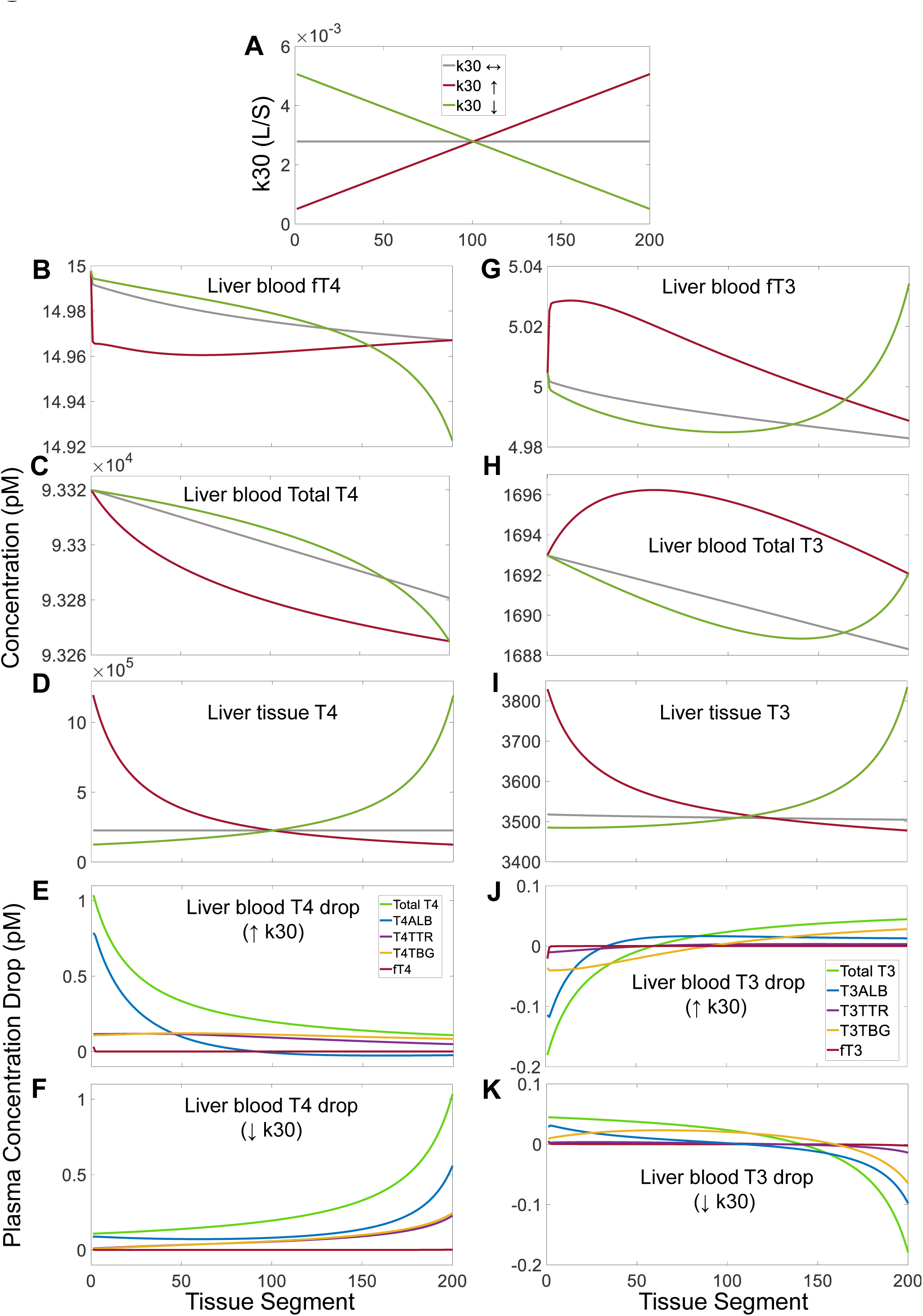
Effects of *k*_30_ gradients across *Liver* segments on TH concentrations in *Liver* and contributions of THBPs to TH loading and unloading in the spatial PBK model. (A) Linearly increasing, decreasing, or constant *k*_30_ gradients implemented as indicated. **(B-D)** T4 concentrations in *Liver blood* and *Liver tissue* as indicated. **(E-F)** Segment-to-segment net differences in plasma concentrations of *fT4*, *T4TBG*, *T4TTR*, *T4ALB*, and *Total T4* as indicated in *Liver blood* for increasing and decreasing *k*_30_ gradients respectively. **(G-I)** T3 concentrations in *Liver blood* and *Liver tissue* as indicated. **(J-K)** Segment-to-segment net differences in plasma concentrations of *fT3*, *T3TBG*, *T3TTR*, *T3ALB*, and *Total T3* as indicated in *Liver blood* for increasing and decreasing *k*_30_ gradients respectively. The same color scheme is used for panels (B-D, G-I) as indicated in (A), for panel (F) as in (E), and for panel (K) as in (J).

When *k*_27_, the rate constant for T3 liver influx, increases or decreases from segment 1 through 200 (Fig. S8A), the gradients of all T4-related variables are barely affected (results not shown). With an increasing *k*_27_ gradient, T3 in *Liver blood* is loaded onto as opposed to unloaded from the THBPs in the upstream quarter of the segments (Fig. S8B). Moving downstream, ALB unloads more T3 than TBG does, but ultimately TBG contribution surpasses ALB contribution. With a decreasing *k*_27_ gradient, ALB initially contributes more T3, while TBG contributes the most T3 throughout the majority of the downstream segments, and all THBPs revert to net loading of T3 toward the venous end (Fig. S8C). The total amount of T3 unloaded by all three THBPs combined changes linearly across the segments (Fig. S8B and S8C). As *k*_27_ varies, *fT3* and *Total T3* in *Liver blood* exhibit nonmonotonic profiles but the percentage changes are negligibly small (Fig. S8D-S8E). T3 in *Liver tissue* is sensitive to *k*_27_ – i.e., as *k*_27_ increases or decreases linearly 10-fold, T3 changes by nearly 10-fold in the same direction (Fig. S8F). When *k*_31_, the rate constant for T3 liver efflux, varies from segment 1 through 200, it produces an effect on T3 (Fig. S9) that is opposite to the effect of varying *k*_27_ but similar to the effect of varying *k*_30_ on T4 (Fig. 11). T3 in *Liver tissue* exhibits a hyperbolic gradient that is sensitive to *k*_27_ (Fig. S9F). Varying *k*_27_ has no effects on T4-related variables (results not shown).

When *k*_26_, the rate constant for T4-to-T3 conversion in *Liver tissue*, varies from segment 1 through 200 by 10-fold (Fig. 12A), the gradients of *fT4* and *Total T4* in *Liver blood* deviate slightly from the case when *k*_26_ is constant (Fig. 12B-12C). When *k*_26_ increases, the gradient of T4 in *Liver tissue* becomes steeper than when *k*_26_ is constant, but the percentage drop remains negligible, only about 0.5% (Fig. 12D, red line). When *k*_26_ decreases, the gradient of T4 in *Liver tissue* is reversed, which increases by about 2% (Fig. 12D, green line). The relative contributions to T4 in *Liver blood* by THBPs are similar to the case when *k*_26_ is constant (Fig. 12E and 12F vs. Fig. 4A). *fT3* and *Total T3* in *Liver blood* exhibit nonmonotonic profiles (Fig. 12G-12H). Despite that the liver T4-to-T3 conversion rate constant *k*_26_ varies by 10-fold across the segments, T3 in *Liver tissue* is insensitive to the changes, increasing or decreasing only by about 3% (Fig. 13I). The changes in the relative contributions of THBPs to T3 unloaded in *Liver blood* (Fig. 12J and 12K) are similar to the case of varying *k*_25_. Varying *k*_34_, the rate constant for T4 metabolism in *Liver tissue*, has a small effect on T4 gradient in *Liver tissue*, which only changes by slightly > 1% from the arterial to venous ends, and the effect on T3 in *Liver tissue* is negligible (Fig. S10). Varying *k*_35_, the rate constant for T3 metabolism in *Liver tissue*, has no effect on T4-related variables (results not shown), and only causes a T3 gradient in *Liver tissue* that varies by < 6% from the arterial to venous ends (Fig. S11).

**Figure 12.**
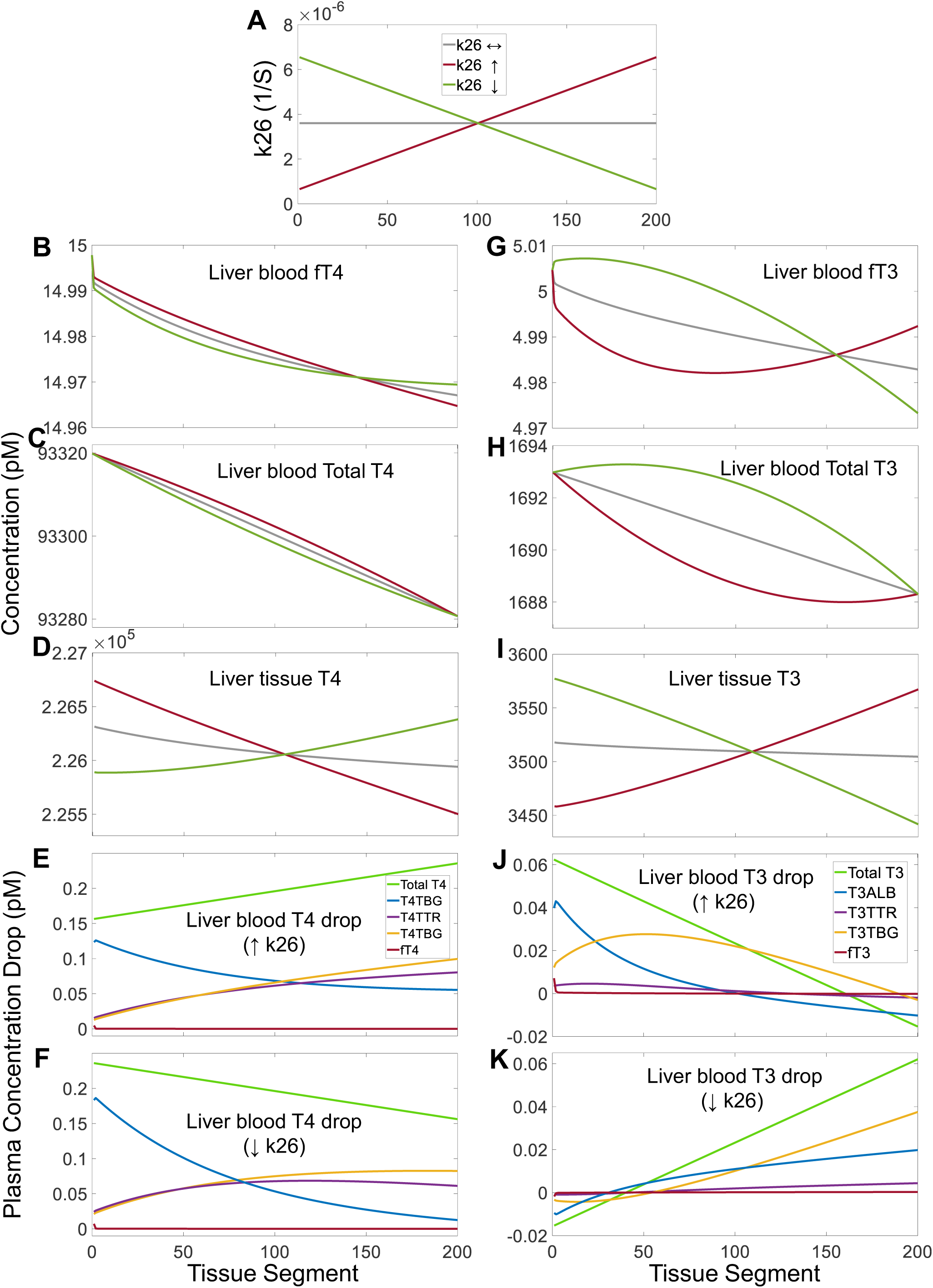
Effects of *k*_26_ gradients across *Liver* segments on TH concentrations in *Liver* and contributions of THBPs to TH loading and unloading in the spatial PBK model. (A) Linearly increasing, decreasing, or constant *k*_26_ gradients implemented as indicated. **(B-D)** T4 concentrations in *Liver blood* and *Liver tissue* as indicated. **(E-F)** Segment-to-segment net differences in plasma concentrations of *fT4*, *T4TBG*, *T4TTR*, *T4ALB*, and *Total T4* as indicated in *Liver blood* for increasing and decreasing *k*_26_ gradients respectively. **(G-I)** T3 concentrations in *Liver blood* and *Liver tissue* as indicated. **(J-K)** Segment-to-segment net differences in plasma concentrations of *fT3*, *T3TBG*, *T3TTR*, *T3ALB*, and *Total T3* as indicated in *Liver blood* for increasing and decreasing *k*_26_ gradients respectively. The same color scheme is used for panels (B-D, G-I) as indicated in (A), for panel (F) as in (E), and for panel (K) as in (J).

Simulations were also conducted for *RB*-related parameters *k*_21_, *k*_28_, *k*_23_, *k*_29_, *k*_24_, *k*_32_, and *k*_33_ respectively (results not shown). In general, when compared to their counterparts in *Liver*, applying gradients to these parameters modulates the THBP contributions to T4 and T3 tissue delivery in a qualitatively similar fashion. Compared to the corresponding parameters in *Liver*, however, T3 in *RB tissue* is much more sensitive to the gradients of *k*_21_, *k*_24_, *k*_28_, *k*_32_, and *k*_33_, but less sensitive to the gradients of *k*_23_ and *k*_29_. T4 in *RB tissue* is still largely insensitive to the gradients of most parameters except *k*_21_ and *k*_28_.

## Discussion

### 1. Contribution of THBPs to TH tissue delivery

#### 1.1 Spatial uniformity of TH tissue delivery and concentration

One of the most prominent functions of THBPs is to transport THs from the thyroid gland through the blood circulation to various target tissues and unload them therein. Our PBK models show minimal drops between the arterial and venous concentrations of free THs (Fig. 3A and 3F, Tables 1 and S1), suggesting that the cells at the vascular entrance of the tissues are exposed to nearly identical TH concentrations as those at the exit. The spatial model further demonstrates that when no parameter gradients exist within the tissue, nearly constant amounts of THs are unloaded in the first through the last tissue blood segments. Removing all three THBPs results in markedly steep drops in the concentrations of *fT4* and *fT3* as well as tissue T4 and T3, which is consistent with the descending TH gradients observed experimentally using *ex vivo* liver perfusion (Weisiger et al. 1986, Mendel et al. 1987, Mendel et al. 1988), and confirms the obligatory role of THBPs in maintaining uniform TH tissue delivery. However, the three THBPs are redundant and thus cross-compensatory in this role, as either one alone appears to be sufficient to maintain relatively constant TH concentrations through tissues (Figs. S2-S4). The sufficiency of ALB alone was previously demonstrated by using *ex vivo* liver perfusion (Mendel et al. 1987). Our simulation further showed that this uniformity may be modulated such that TH tissue delivery and concentrations become graded when there exist expression or activity gradients of the membrane TH transporters and certain metabolic enzymes within the tissue.

#### 1.2 Contribution of individual THBP species

It has been argued that ALB supplies the most THs to tissues due to THs having the shortest residence time on ALB; in contrast a smaller fraction of THs is supplied by TBG since THs have much longer residence time on TBG (Mendel 1989, Schussler 2000). It has also been proposed that TTR makes the most contribution to immediate T4 tissue delivery because of its Goldilocks properties – it has the intermediate binding affinity for T4 and the abundance of T4TTR is intermediate among the three T4-THBPs (Robbins 2002, Richardson 2009, Alshehri et al. 2015). The roles of TTR and TBG were also compared analogously to bank accounts, where TTR is the checking and TBG the saving accounts such that T4TTR is tapped first for T4 tissue delivery and then it is replenished by T4TBG later on after they leave the tissue (Robbins 2002, Richardson 2009, Hamers et al. 2020). However, our models suggest neither of the two hypotheses on the roles of ALB and TTR relative to TBG is valid.

Simulations of both the nonspatial and spatial models indicate that TBG, TTR and ALB contribute comparable amounts of T4 in *Liver*, but TBG plays a dominant role in *RB* (Fig. 2 and Tables 1 and S1). This is because the amounts of T4 unloaded by T4-THBPs are determined by both their abundances and T4 residence time relative to the blood tissue transit time. TBG binds the most T4 (75%) out of the three THBPs, which compensates for the longer residence time and thus slower release of T4. The longer blood transit time in *RB* provides more time for TBG to unload T4, which explains the much higher contribution of TBG to tissue T4 in *RB* than in *Liver*. Therefore, for slowly perfused tissues, the contribution to tissue T4 follows the order of the abundances of T4-THBPs. The residence time of T3 on THBPs are generally much shorter than T4, especially for T3TBG and T3TTR, therefore the unloaded amounts are more aligned with their relative abundance rather than the residence time. This explains the consistent order of contribution to T3 by THBPs in both tissue compartments (Fig. 2), where TBG contributes about 3 times more than ALB, and ALB nearly 4 times more than TTR.

Generally, for tissue uptake of a substance bound to plasma proteins, the dissociation rate, rate of transportation or diffusion into the tissues, and blood perfusion rate can all play some limiting roles depending on the parameter conditions (Weisiger 1985). The dissociation of THs from a THBP species could be rate-limiting in providing THs for tissue uptake, the significance of which depends on the tissue blood perfusion rate. For very slowly perfused tissues, the THBP contribution to tissue TH follows the order or even the proportion of the abundances of the TH-THBPs as demonstrated by our simulations when the RB flow rate is decreased by 4-fold from default value (Fig. 9C, 9D, and 9G). In contrast, for rapidly perfused tissues, the percentage contribution by ALB will become higher while that by TBG will become lower since the instantaneous T4 release matters more (Fig. 9A and 9E). Ultimately it is the absolute dissociation rate, which is determined by both the TH residence time on THBP and the abundance of TH-THBP, that dictates the amount of TH released in the most rapidly perfused tissues. Unlike in tissues that are T3 sinks, in T3-source tissues where T3 is produced from T4 locally with no or little metabolic removal, there will be net loading of the THBPs by T3 in the tissue blood. However, our simulation showed that in the latter case the amount of T3 loaded still follows the order of T3-THBP abundances regardless the tissue is fast or slowly perfused (Fig. 9F and 9H).

In extremely rapidly perfused extrathyroidal tissues, it is conceivable that ALB is the most liquid checking account, from which the vast majority of T4 and T3 are withdrawn and delivered to the tissue. As the blood goes through the thyroid, ALB is quickly loaded with nearly half of the newly synthesized T4, while TBG and TTR are loaded with the remaining half (Fig. 2A, Tables 1 and S1). Then as the thyroid venous blood mixes with the venous blood coming back from other tissues, T4 will be transferred from ALB to TBG and TTR, as demonstrated by the positive or negative sign of the difference between the association and dissociation rates for each TH/THBP pair in *Body Blood* (Table S2). For T3, because of the relatively faster binding kinetics and lower binding affinities between T3 and THBPs compared with T4, TBG seems to be the THBP that is loaded with the most T3 in *Thyroid blood* (Fig. 2B, Tables 1 and S1). Interestingly, after mixing with the venous blood coming back from other tissues in *Body Blood*, both ALB and TTR transfer T3 to TBG (Table S2).

If T4TBG is too slow to release enough T4, it begs the question why TBG alone is sufficient to unload a similar amount of T4 in *Liver* and *RB* as when all three THBPs are present (Figs. S2 vs. 3). This is because with TBG alone, if *T4TBG* does not unload enough T4 in the tissue, *T4TBG* in the venous blood would initially leave the tissue at a higher concentration than it would be when T4 is sufficiently unloaded. After perfusing through the thyroid, *T4TBG* will be recharged by the secreted T4 to an even higher level. With *T4TBG* in the arterial blood coming back at a higher level to the tissue, the dissociation rate will be higher, thus more T4 is unloaded. The slightly higher arterial *T4TBG* concentration is evidenced by the levels shown for segment 0 in Fig. S2 compared with Fig. 3. In addition, because of the difference in residence time, a larger drop of free THs in the first segment occurs when TBG is the only THBP (Fig. S2A and S2C), which contrasts with the smaller drop of free THs when ALB is the only THBP (Fig. S4A and S4C). The initial larger drop in free THs moves TH-TBG further away from equilibrium, reducing the association rate and allowing sufficient amounts of THs to be released from TH-TBG.

#### 1.3 Spatially dependent variation of THBP contribution

An interesting and novel finding of the present study is that the three THBPs unload THs in a spatially dependent manner, where ALB generally contributes the most at the beginning whereas TBG contributes the most at the end of the perfused tissue. This is due to different residence times among the THBPs relative to the tissue blood transit time and different abundances of TH-THBPs. At the beginning of tissue perfusion, there is not enough time for THs to dissociate from TBG or TTR due to their longer residence time. As a result, ALB, which has a shorter residence time, will unload more. The relative contribution of the THBPs to tissue THs in the most upstream segments thus follows the order of the absolute dissociation rates of each TH-THBP (Figs. 5 and 6). As the blood continues to move through the tissue with free THs dropping, TBG and TTG will become further removed from their equilibrium, resulting in a higher differential between the dissociation rate and association rate, thus more THs are unloaded from TBG and TTR. Therefore, with longer perfusion time, the TH contribution by TBG and TTR will catch up, while that by ALB will lessen. If the perfusion time is long enough, like in *RB* for T3, the distances from equilibrium for the three THBPs will approach some constants (Fig. 6D-6F), and the relative contribution by these three TH-THBPs will reach a fixed ratio proportional to their abundances (Fig. 4D). At such a state, TBG is the one that is mostly removed from equilibrium, while ALB is closest to equilibrium, with TTR in the middle. Being the furthest removed from equilibrium allows TBG to deliver the most TH despite its tight grip on T4 and T3. For tissues acting as a T3 source, the loading of T3 to the three THBPs follow the same trend of spatial dependence as the case of unloading. Lastly, the spatially dependent, nonlinear kinetics of THBPs may explain some of the discrepancies in the overall amounts of unloaded THs in *Liver* predicted by the spatial vs. nonspatial models. The spatial model predicts that T4ALB unloads a slightly higher amount of T4 than T4TBG does in *Liver* (Fig. 2A and Table 1), while the nonspatial model predicts the opposite (Table S1). Such discrepancy underscores the importance of modeling tissue blood flow in greater details for more accurate predictions.

The above conclusions are based on simulations with the assumption that the influx, efflux, and metabolism parameter values are constant across the tissue segments. Our simulations with spatial parameter heterogeneities indicate that they tend to modulate the contributions of different THBP species to TH tissue delivery and can ultimately impact the heterogeneity of intra-tissue concentrations of T4 and T3 in some cases. Because of the near-equilibrium kinetics of TH influx and efflux between the blood and tissues, especially for T4 in *Liver* and *RB* and for T3 in *Liver* (Bagga et al. 2023), the intra-tissue T4 and T3 concentrations are mostly sensitive to gradients of the respective influx and efflux parameters. In contrast, they are much less sensitive to gradients of metabolism parameters. For tissues operating in non-equilibrium mode of TH exchange, the tissue T4 and T3 concentrations are sensitive to gradients of metabolism parameters. These properties of local TH gradient regulation appear to be utilized for pattern formation during development. One example is how T3 determines the pattern of the retina. In mice, T3 concentration forms a roughly 2.5-fold dorsal-to-ventral decreasing gradient in the nascent retina on postnatal day 10 (Roberts et al. 2006). This gradient drives the formation of a pattern where the medium-wavelength opsin expressing cone cells dominate in the dorsal retina while the short-wavelength opsin expressing cone cells dominate in the ventral retina. The T3 gradient was not due to variations of DIO expression because DIO1 and 2 levels were undetectable while DIO3 displayed a uniform distribution in the developing retina. It was argued that graded TH transporter expression may create the T3 gradient in the retina (Roberts et al. 2006), which is consistent with our findings that intra-tissue T3 distribution is sensitive to influx and efflux parameter gradients. In comparison, during metamorphosis of the frog, there is a DIO3 gradient concentrated in the dorsal half of the developing retina, which leads to T3 accumulation in the ventral half that drives the proliferation of ventral ciliary marginal zone (CMZ) cells and the formation of the ipsilateral projection (Marsh-Armstrong et al. 1999). This sensitivity of T3 accumulation to DIO3 gradient suggests that the developing retina of the frog may not operate in equilibrium mode for blood-tissue exchange of THs.

The liver lobule is known for its zonal heterogeneity in gene expression (Gebhardt and Matz-Soja 2014, Halpern et al. 2017). To meet different functional demands for THs in different zones, there may exist spatial gradients of transporters and enzymes in their abundance and affinity such that the rates of local influx, efflux, and metabolism of THs, and consequently, the hepatic TH concentrations and their effects may vary across zones. Based on our analysis, even if there are gradients in the abundance and/or activities of the metabolic enzymes, as long as there is no gradients for the transporters, T4 and T3 are not expected to form tangible gradients within the liver lobule. By perfusing the rat liver through the portal vein with human serum containing ^125^I-labeled T4, it was shown that the hepatic T4 deposition was similar in the different layers of the rat liver lobule (Mendel et al. 1987), suggesting that there is no heterogeneous distribution of T4 transporters. Antegrade and retrograde liver perfusions in the absence of THBPs further corroborated this by showing that the periportal-to-central capacities for T4 uptake are relatively constant, at least in rat livers (Weisiger et al. 1986, Mendel et al. 1988). Since there is no T3 gradient in the lobule, any heterogeneity in T3 effects will likely result from zonal distribution of thyroid receptors or downstream effector proteins.

### 2. Evolutionary perspective on THBPs and their biological functions

Power et al. reviewed a wide array of studies on primitive vertebrates including fish, amphibians, reptiles, and birds, suggesting that ALB emerged earlier than TTR evolutionarily (Power et al. 2000). Since ALB binds many compounds weakly, it is not surprising that more TH-specific THBPs arose as species evolved. TBG appeared the latest with mammals being the only animal class that have all three THBPs (McLean et al. 2017). Interestingly, these THBPs share some homologous sequences in the domains involved in TH binding, especially a 5-residue TH-binding motif, with TH-binding apolipoproteins, cell membrane TH transporters, and TH receptors, suggesting the origin of a common ancestor gene (Benvenga and Guarneri 2018).

Our simulations indicate that not all three THBP species are needed to maintain normal free TH concentrations and uptake within a tissue, which is consistent with a number of experimental and clinical studies. For instance, T4 uptake in the liver, kidney and brain of the Nagase analbuminemic rats was not different from normal rats (Mendel et al. 1989). Adult TTR knockout mice are euthyroid with normal circulating free T4, T3, and TSH but lower total T4 and T3 concentrations (Episkopou et al. 1993, Palha et al. 1994). TBG-deficient patients have similar TH profiles and are also euthyroid (Mimoto and Refetoff 2020).

If only one or two of the three THBP species is sufficient, it thus follows that the THBPs that emerged later may have additional functions besides supplying THs to tissues. TTR can be synthesized in the choroid plexus in the brain and placenta where it plays a unique role in transporting THs against the local concentration gradients (Landers et al. 2013, Richardson et al. 2015, Rabah et al. 2019). Although the adult TTR knockout mice are euthyroid, postnatal mice exhibited delayed weaning and development of the central nervous system, bones, intestine, and muscle (Monk et al. 2013). The late-emerging TBG appears to provide several additional functions. T4 and T3 are predominantly associated with TBG in the plasma, and moreover, among the three THBPs, TBG has the highest binding affinities for T4 and T3. Therefore, the buffering function of THBPs against transient TH perturbations is mainly carried out by TBG. Furthermore, the total amount of T4 in the plasma is equivalent to about 2-days’ worth of T4 produced by the thyroid; therefore, TBG may function as an extrathyroidal reservoir of THs, in addition to the intrathyroidal store in thyroglobulin, in the event of disruption of TH production. The relative contribution of THBPs to tissue THs does not appear to be fixed and may vary under different physiological and pathological conditions. For instance, TBG can be regulated during acute inflammation, where locally released serine proteases can cleave TBG to a low-binding-affinity form that can unload more THs to the inflammatory sites (Pemberton et al. 1988, Jirasakuldech et al. 2000). During pregnancy, the maternal TBG level steadily doubles through the mid-term because of the rising estrogen and reduced plasma clearance, which may function to titrate the increasingly produced maternal THs and prevent them from rising too high (Moleti et al. 2014).

### 3. Limitations, potential applications, and future directions

Our current model does not include the HPT negative feedback loop. This is because we are studying the local intra-tissue TH and THBP kinetics at steady state with constant inputs of these species from the arterial blood. Therefore, it is not necessary to consider any actions of THs on TSH and vice versa, the inclusion of which in the model is not expected to change the results and conclusions in the present study. The uncertainty in the amounts of THs unloaded in the tissue blood by each THBP species predicted by our models lies in the values reported in the literature for the relative abundance of TH-THBPs and dissociation rate constants, which can vary depending on individuals and their physiological conditions as well as the experimental methodologies. The fraction of T4TBG may be overstated when measured with zone electrophoresis by as much as 10% because of the comigration of T4-binding high-density lipoprotein (HDL) and some serpin proteins with TBG (Benvenga et al. 2002). Despite such overestimation, the abundance of T4TBG is still expected to be much higher than T4TTR and T4ALB. Therefore, for slowly perfused tissues, the order of contribution to T4 by each THBP species would remain largely unchanged because it primarily scales with the abundance of each TH-THBP species. In contrast, the predicted relative contribution by each THBP species for the liver, a rapidly perfused tissue, is more uncertain because both the abundances and dissociation rate constants of THBPs play a role here, but fewer experiments have been done to determine the latter parameters.

While the three THBPs modeled here play major roles in regulating the amounts of THs unloaded and thus available for tissue uptake, previous research indicated that lipoproteins including HDL and low-density lipoprotein (LDL) also affect TH influx into and efflux out of cells (Benvenga and Robbins 1990, Benvenga and Robbins 1998). In static cultures of skin fibroblasts or hepatocytes, the net efflux of T4 out of the cultured cells was facilitated by the lipoproteins as well as by THBPs (Benvenga and Robbins 1998). The facilitation can be partially explained by these proteins acting as a sink to lower free T4 in the medium to reduce re-entry. As a matter of fact, as long as the free fraction of T4 was titrated to the same level, regardless of using either TBG, TTR, or ALB in the cell culture, T4 influx or uptake by the cells was reduced to a similar magnitude (Benvenga and Robbins 1990). On an equal molar basis, the order of efflux facilitation was TBG > LDL > TTR > HDL >> ALB, which cannot be explained by the relative binding affinities of these proteins for T4 because lipoproteins are generally weaker T4 binders. It was postulated that LDL and HDL may engage some nonspecific physical contact with the plasma membrane and their hydrophobic nature helps to desorb T4 off of the plasma membrane. LDL is better than HDL in facilitating the efflux because of its higher lipophilic content. Counterintuitively, it was also shown that LDL may facilitate T4 uptake (Benvenga and Robbins 1990). This occurs in cells expressing LDL receptor (LDLR) (Benvenga and Robbins 1990), where after T4-carrying LDL docks to LDLR, T4 can be released and internalized by cells. Such LDLR-mediated T4 entry may play a role in regulating lipid metabolism in local tissues. Future iterations of our model may consider including LDL and HDL-mediated TH tissue uptake.

The liver is an organ that can take up and secrete fatty acids, which can compete with THs for ALB binding. Serum fatty acid concentrations can fluctuate as the blood transits through different segments of the liver tissue when they are taken up from or secreted into the blood. Thus, theoretically it is possible that such fluctuations may impact the local TH binding to ALB and affect the spatial model accuracy. But the impact is unlikely to be significant, because at physiological concentrations, THs only occupy a tiny fraction (about 0.0016%) of the binding sites on ALB (Schussler 2000), so there are likely plenty sites available for fatty acids and other molecules such as steroid hormones without tangibly displacing THs from ALB. Moreover, experimental studies demonstrated that when myristate, a fatty acid that can bind to ALB with high affinity (Fujiwara and Amisaki 2008), is present at concentrations 30-fold greater than physiological levels, human serum ALB still retains a high-affinity binding site for T4 (the affinity was only reduced by half), due to some conformational changes of ALB (Petitpas et al. 2003). The intracellular distribution of THs can be heterogeneous as well. For instance, the nucleus tends to accumulate much higher free T3 than the cytosol (Oppenheimer and Schwartz 1985). Future iterations of the model may consider further compartmentalizing a tissue into cytosolic and nuclear spaces if needed.

Because pertinent parameter values for modeling THs in specific individual tissues are generally not available, here all extrahepatic and extrathyroidal tissues had to be modeled as a single compartment *RB* whose behavior may represent the average of these tissues. An individual tissue can be different than the average in their blood perfusion rate, TH uptake rate, and the local production and metabolism rates depending on the expression/activity levels of DIOs (Goemann et al. 2018). Future updates to the model could split *RB* into separate tissue compartments as sufficient tissue-specific data such as TH tissue concentrations, transporter and TH-metabolizing enzyme related kinetic parameters become available.

For simplicity, we used a linear gradient and varied one parameter at a time to examine their impact on THBP contributions to TH tissue delivery and tissue TH concentrations. In reality, the gradients of transporters and metabolic enzymes can vary in shape which could be exponential, nonmonotonic, or others. Moreover, the gradients of THs may not be controlled by a single protein or enzyme. Rather, their gradients may multiplex to collectively regulate local TH distributions within a tissue. As more tissue-specific spatial information becomes available, they can be included in future models.

Our models and their future iterations can have several potential applications. It is well known that the thyroid system is subjected to circadian regulation. In humans there are strong daily rhythms of TSH and fT3 accompanied with less pronounced rhythm of fT4 (Nimalasuriya et al. 1986, Russell et al. 2008). A strong temporal relationship between TSH and fT3 has been suggested to be associated with familial longevity (Jansen et al. 2015). Since THBPs carry THs in blood circulations, they may affect the relative timing of TH tissue distribution. Applying control theory, we are currently using the PBK model to study the amplitude and phase relationships between TSH, fT3, fT4 and tissue THs and the potential mechanisms underpinning the circadian patterns. Our model may also help with clinical chronotherapy to guide the timing of thyroid drug administration to ensure that the fluctuations of plasma THs are more closely aligned with the natural circadian rhythm. Many environmental chemicals can interfere with TH binding to THBPs because of their similar structures to THs. These endocrine-disrupting chemicals (EDCs) include polychlorinated biphenyls (PCBs), dibenzo-p-dioxins, dibenzofurans, and brominated flame retardants (Cheek et al. 1999, Hamers et al. 2006). EDCs, especially those with high binding affinities, can displace THs from THBPs to cause a potential increase in fT4 or fT3 levels (Brouwer et al. 1998, Noyes et al. 2019). Using the nonspatial version of the PBK model, we have recently predicted that for persistent exposure to these EDCs the impact on free THs is minimal, whereas daily intermittent exposure to EDCs with short half-lives may cause concern as free THs can fluctuate to an extent that interferes with the circadian rhythm (Bagga et al. 2023). The spatial model can be used to further investigate the impacts of thyroid EDCs on tissue TH delivery by THBPs.

## Conclusion

In summary, we have constructed a spatial human PBK model with explicit considerations of TH and THBP interactions and explored their differential contributions in supplying TH to tissues. While some of the predictions remain to be experimentally validated, the model provides novel insights into the debated roles of THBPs in this regard and local TH regulations.

## Funding Acknowledgements

This research was supported in part by NIEHS Superfund Research grant P42ES04911 and NIEHS HERCULES grant P30ES019776.

## Author Contributions

QZ conceived the model structure, ADB constructed and simulated the model in MATLAB, ADB and QZ conducted the parameter justification, parameter estimation, and formal analysis of simulation results, ADB and QZ wrote the initial draft and revised the manuscript. BPJ critically reviewed and revised the manuscript.

## Data Availability Statement

All MATLAB code will be available at https://github.com/pulsatility/2024-TH-PBK-Model.git.

