## Supplemental Figures for "Spatially Dependent Tissue Distribution of Thyroid Hormones by Plasma Thyroid Hormone Binding Proteins"

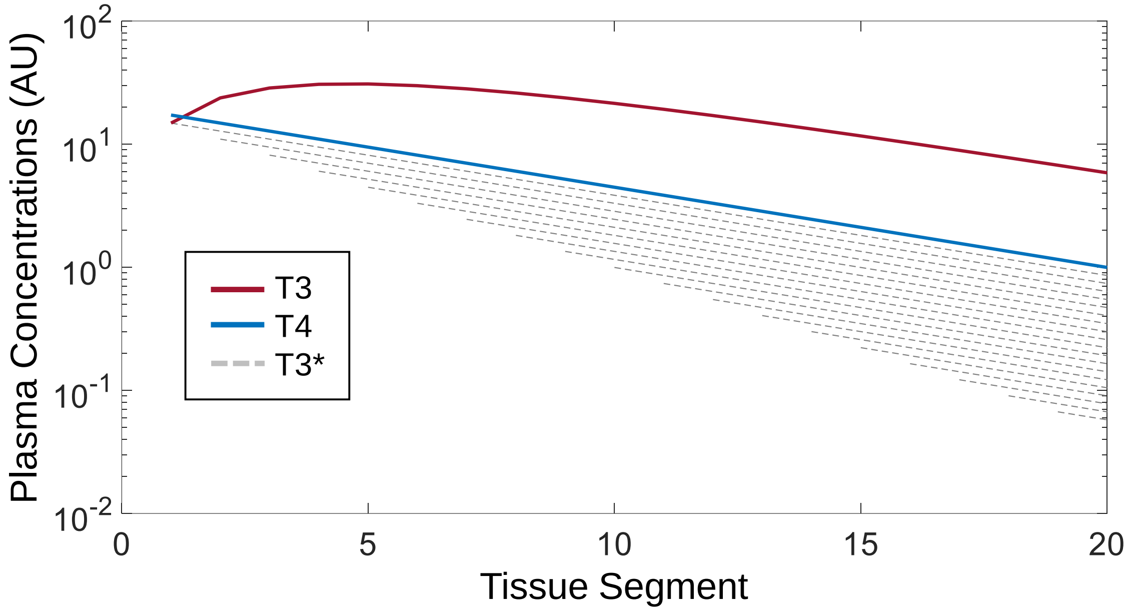


**Figure S1. Illustration of the origin of the nonmonotonic T3 gradient in tissue blood in the absence of all three THBPs by a simple mathematical model.** In the absence of any THBPs, T3 produced from T4 in each tissue segment will leave the tissue segment into the corresponding blood segment and flow with the blood to downstream segments, forming an exponentially descending gradient of its own (dashed gray lines) as it is taken up by downstream segments and metabolized, as T4 does (blue line). The total T3 abundance in each segment will be proportional to the sum of T3 produced *de novo* from T4 in that segment, T3 produced from T4 in all upstream segments flowing downstream, and T3 coming in from the arterial blood which also decays exponentially. The further downstream a segment, the greater number of upstream segments from which it will receive T4-converted T3. In the meantime, however, the T3 coming from these upstream segments will arrive at a lower concentration as it is distributed to the interim tissue segments. These two opposing forces result in a nonmonotonic T3 gradient (maroon line), which is the sum across all the dashed gray lines. The model in MATLAB is provided at the GitHub site associated with the paper. 20 segments are used for uncluttered visualization.

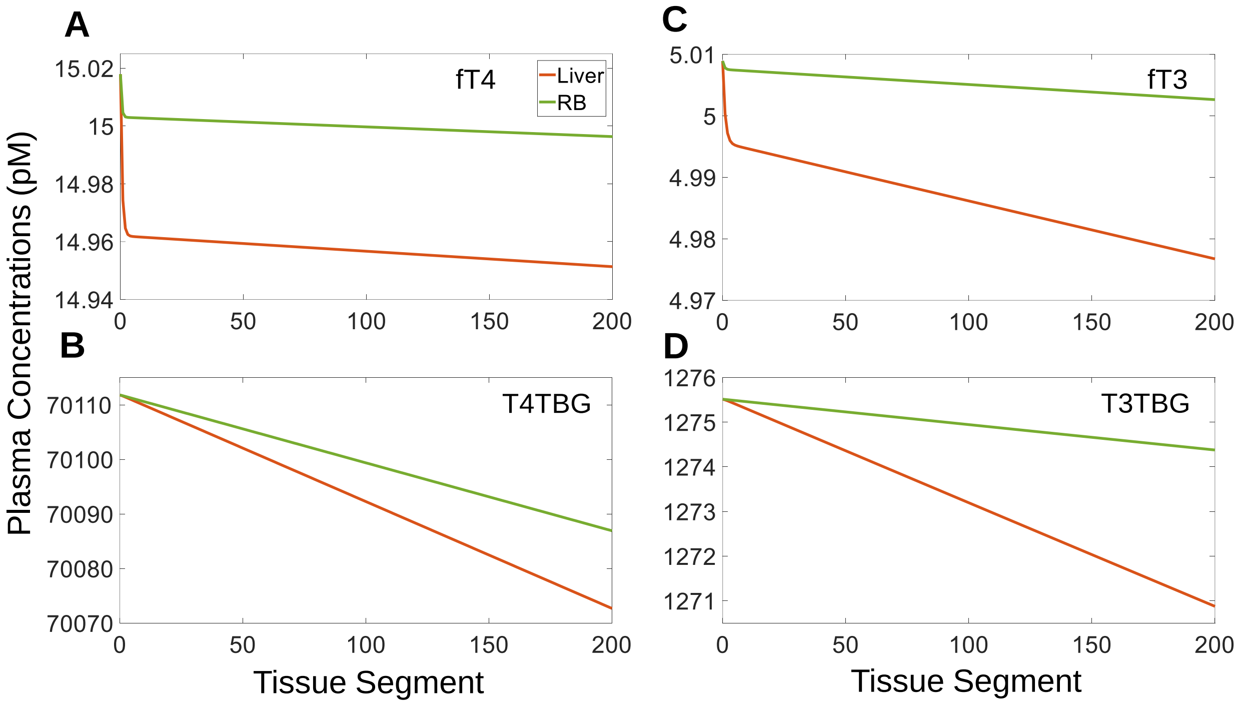


**Figure S2. Plasma concentration gradients of free THs and TH-TBGs in tissue blood with TBG as the only THBP present in the spatial PBK model. (A-B)** Plasma concentrations of *fT4* and *T4TBG* in *Liver blood* (orange) and *RB blood* (green) respectively. **(C-D)** Plasma concentrations of *fT3* and *T3TBG* in *Liver blood* (orange) and *RB blood* (green) respectively. Concentrations in segment 0 represent the plasma concentrations in arterial blood.

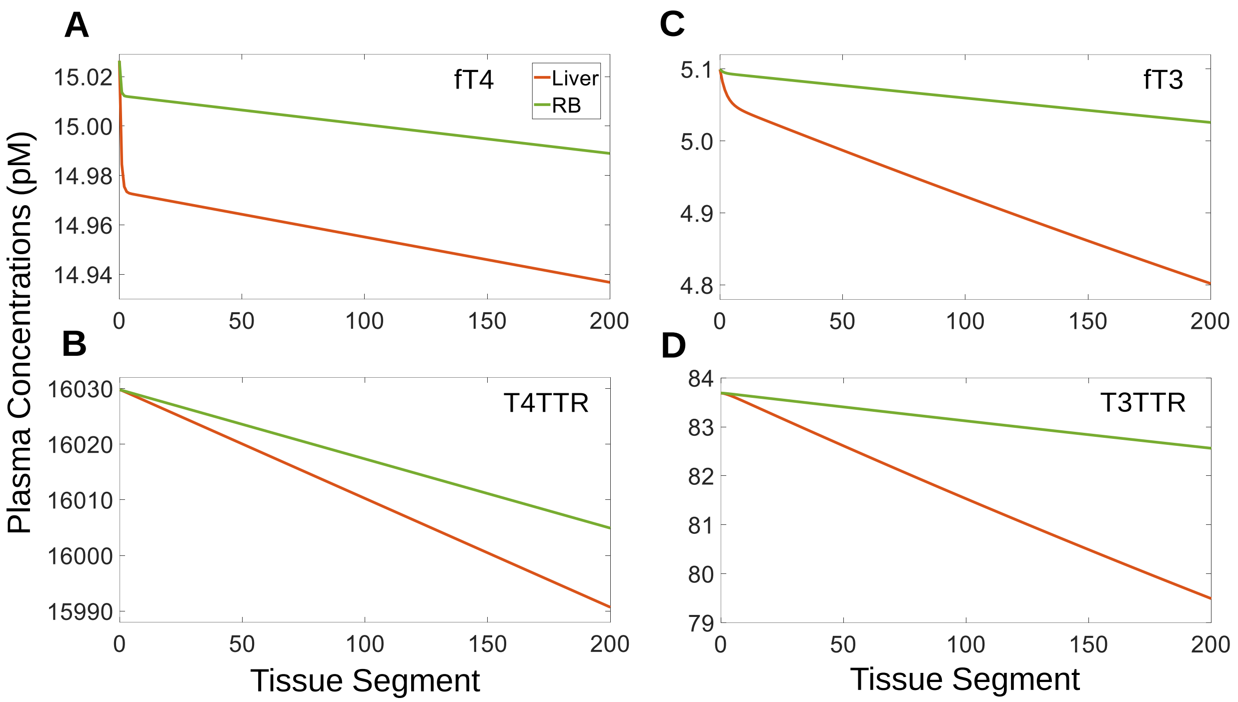


**Figure S3. Plasma concentration gradients of free THs and TH-TTRs in tissue blood with <u>TTR as the only THBP present</u> in the spatial PBK model. (A-B)** Plasma concentrations of *fT4* and *T4TTR* in *Liver blood* (orange) and *RB blood* (green) respectively. **(C-D)** Plasma concentrations of *fT3* and *T3TTR* in in *Liver blood* (orange) and *RB blood* (green) respectively. Concentrations in segment 0 represent the plasma concentrations in arterial blood.

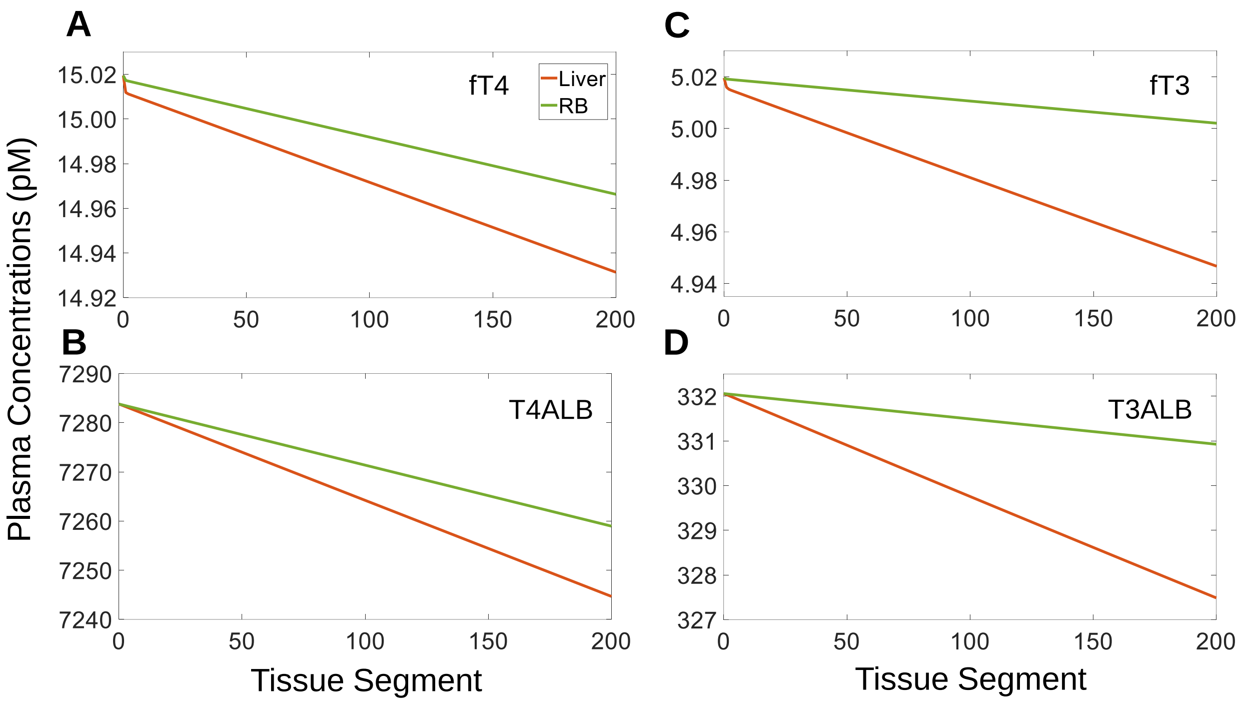


**Figure S4. Plasma concentration gradients of free THs and TH-ALBs in tissue blood with <u>ALB as the only THBP present</u> in the spatial PBK model. (A-B)** Plasma concentrations of *fT4* and *T4ALB* in *Liver Blood* (orange) and *RB Blood* (green) respectively. **(C-D)** Plasma concentrations of *fT3* and *T3ALB* in *Liver blood* (orange) and *RB blood* (green) respectively. Concentrations in segment 0 represent the plasma concentrations in arterial blood.

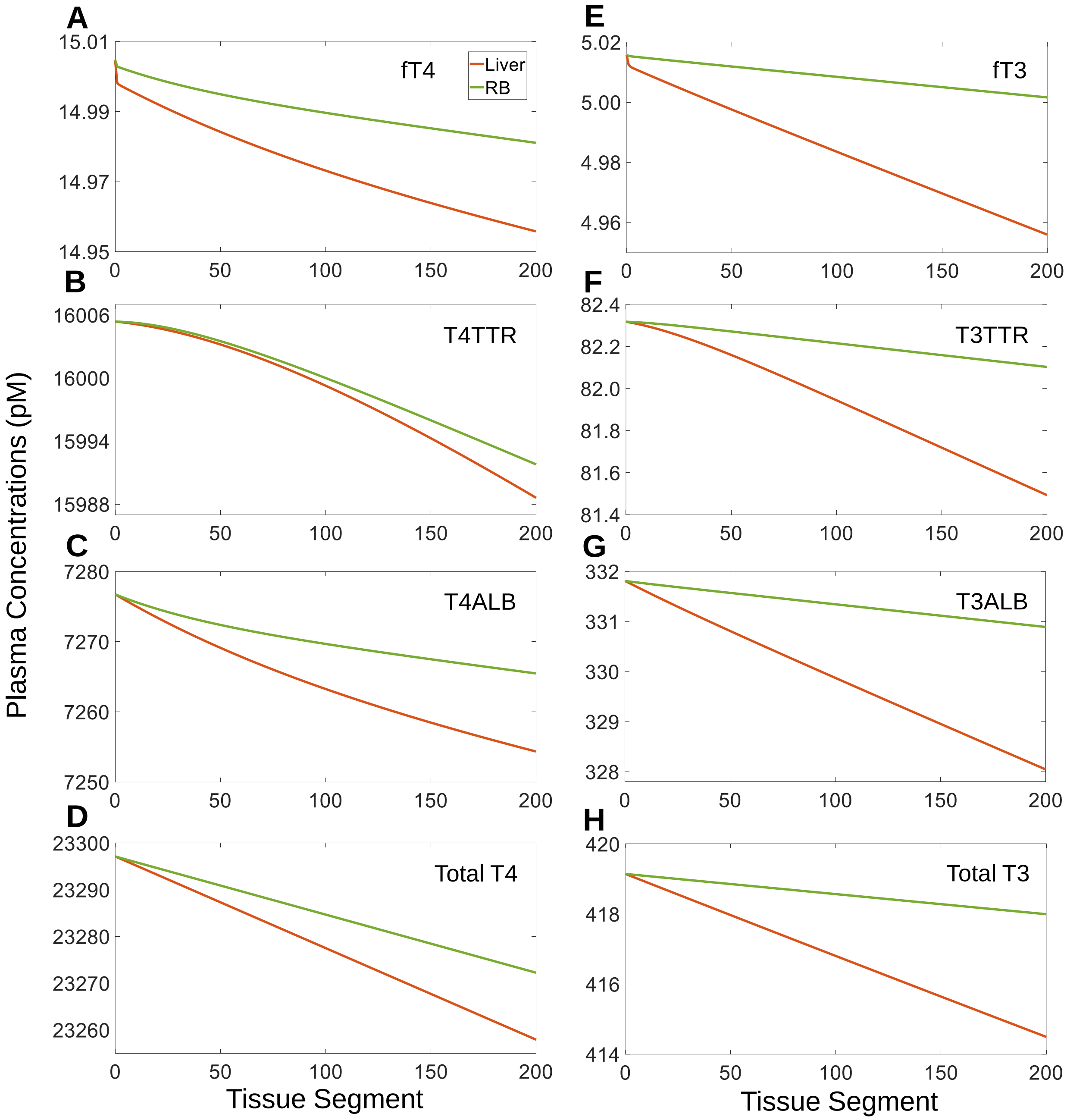


**Figure S5. Plasma concentration gradients of free THs and TH-THBPs in tissue blood <u>in the absence of TBG</u> in the spatial PBK model. (A-D)** Plasma concentrations of *fT4*, *T4TTR*, *T4ALB*, and *Total T4* in *Liver blood* (orange) and *RB blood* (green) respectively. **(E-H)** Plasma concentrations of *fT3*, *T3TTR*, *T3ALB*, and *Total T3* in *Liver blood* (orange) and *RB blood* (green) respectively. Concentrations in segment 0 represent the plasma concentrations in arterial blood.

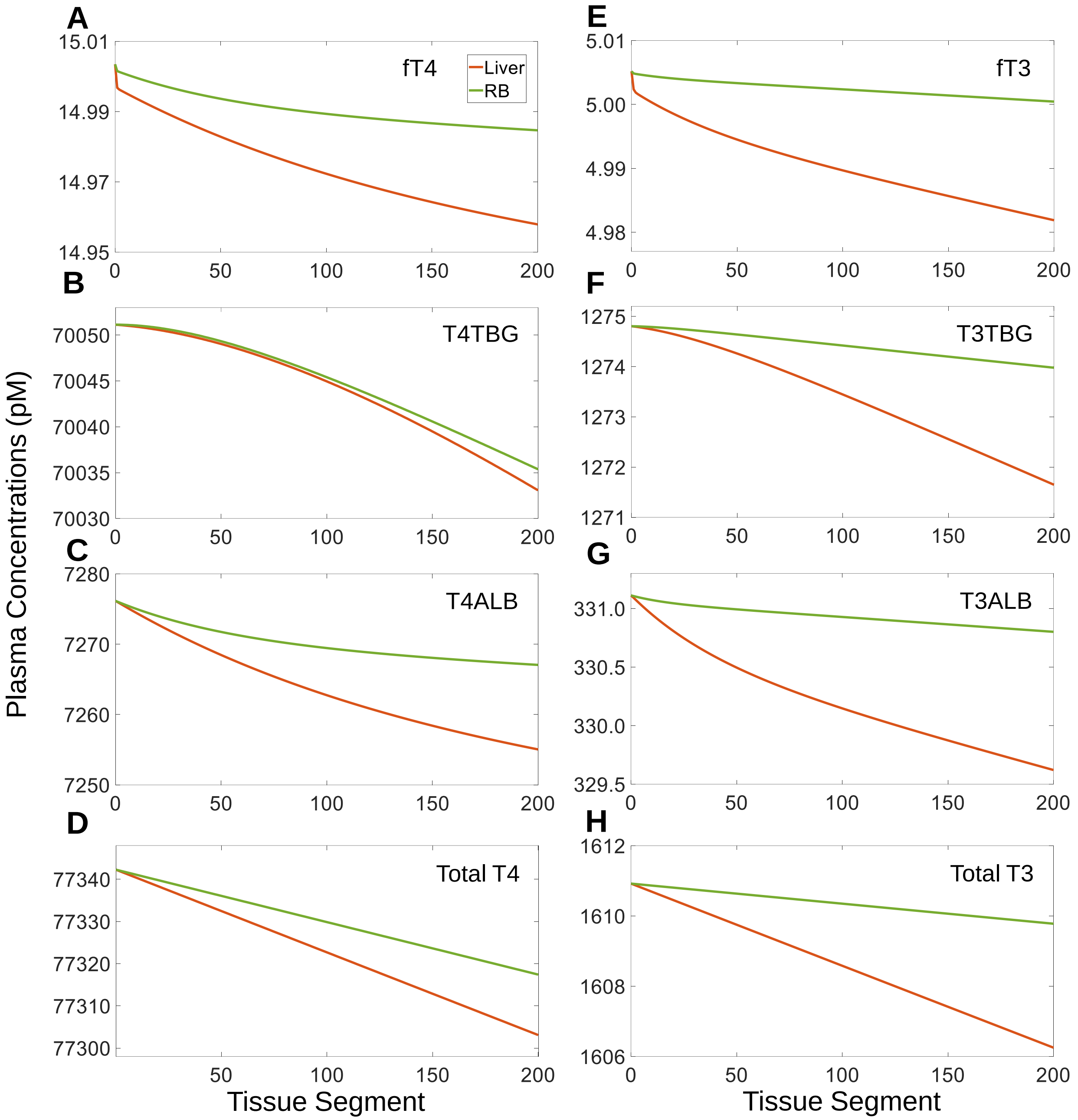


**Figure S6. Plasma concentration gradients of free THs and TH-THBPs in tissue blood <u>in the absence of TTR</u> in the spatial PBK model. (A-D)** Plasma concentrations of *fT4*, *T4TBG*, *T4ALB*, and *Total T4* in *Liver blood* (orange) and *RB blood* (green) respectively. **(E-H)** Plasma concentrations of *fT3*, *T3TBG*, *T3ALB*, and *Total T3* in *Liver blood* (orange) and *RB blood* (green) respectively. Concentrations in segment 0 represent the plasma concentrations in arterial blood.

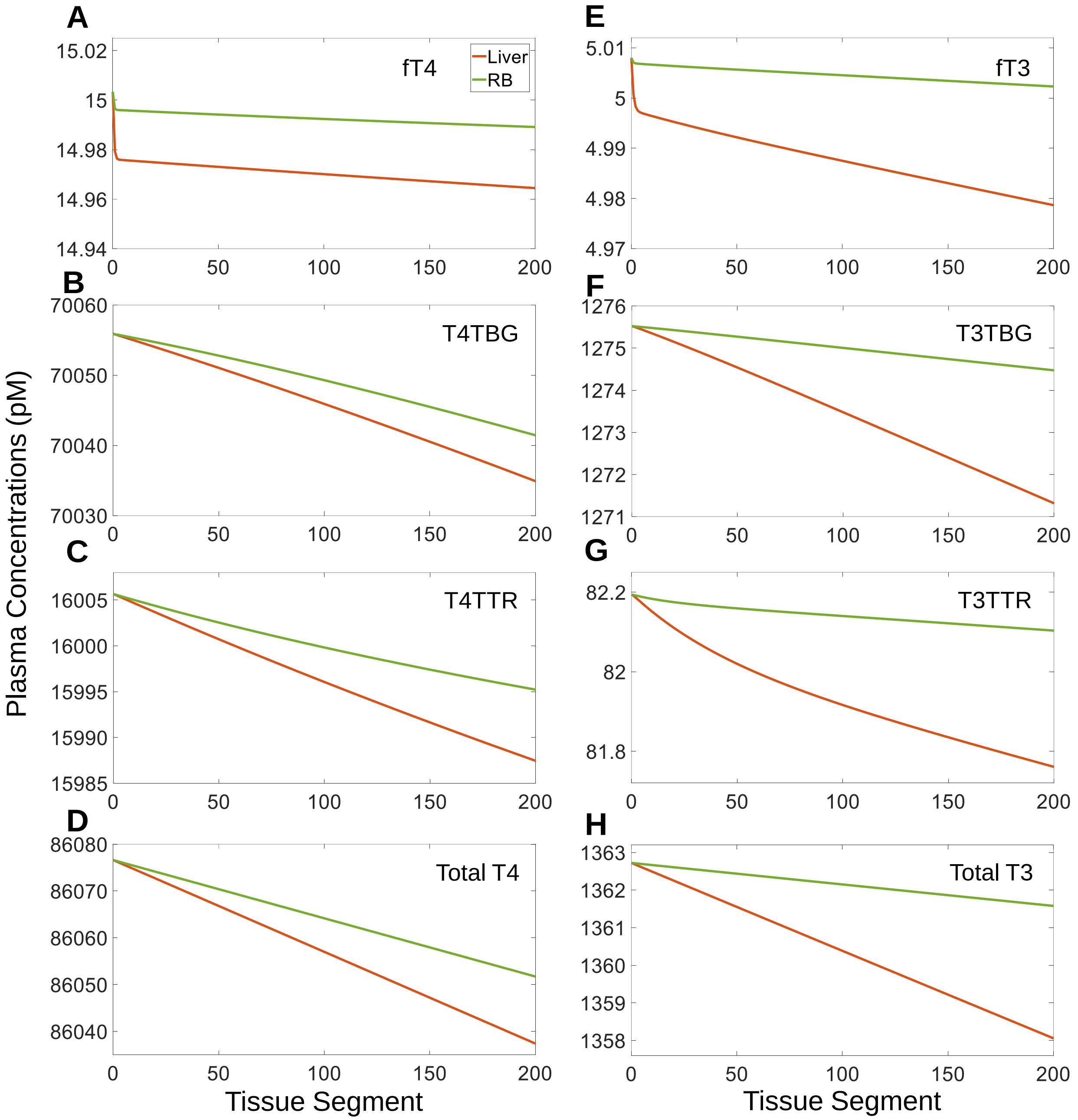


**Figure S7. Plasma concentration gradients of free THs and TH-THBPs in tissue blood <u>in the absence of ALB</u> in the spatial PBK model. (A-D)** Plasma concentrations of *fT4*, *T4TBG*, *T4TTR*, and *Total T4* in *Liver blood* (orange) and *RB blood* (green) respectively. **(E-H)** Plasma concentrations of *fT3*, *T3TBG*, *T3TTR*, and *Total T3* in *Liver blood* (orange) and *RB blood* (green) respectively. Concentrations in segment 0 represent the plasma concentrations in arterial blood.

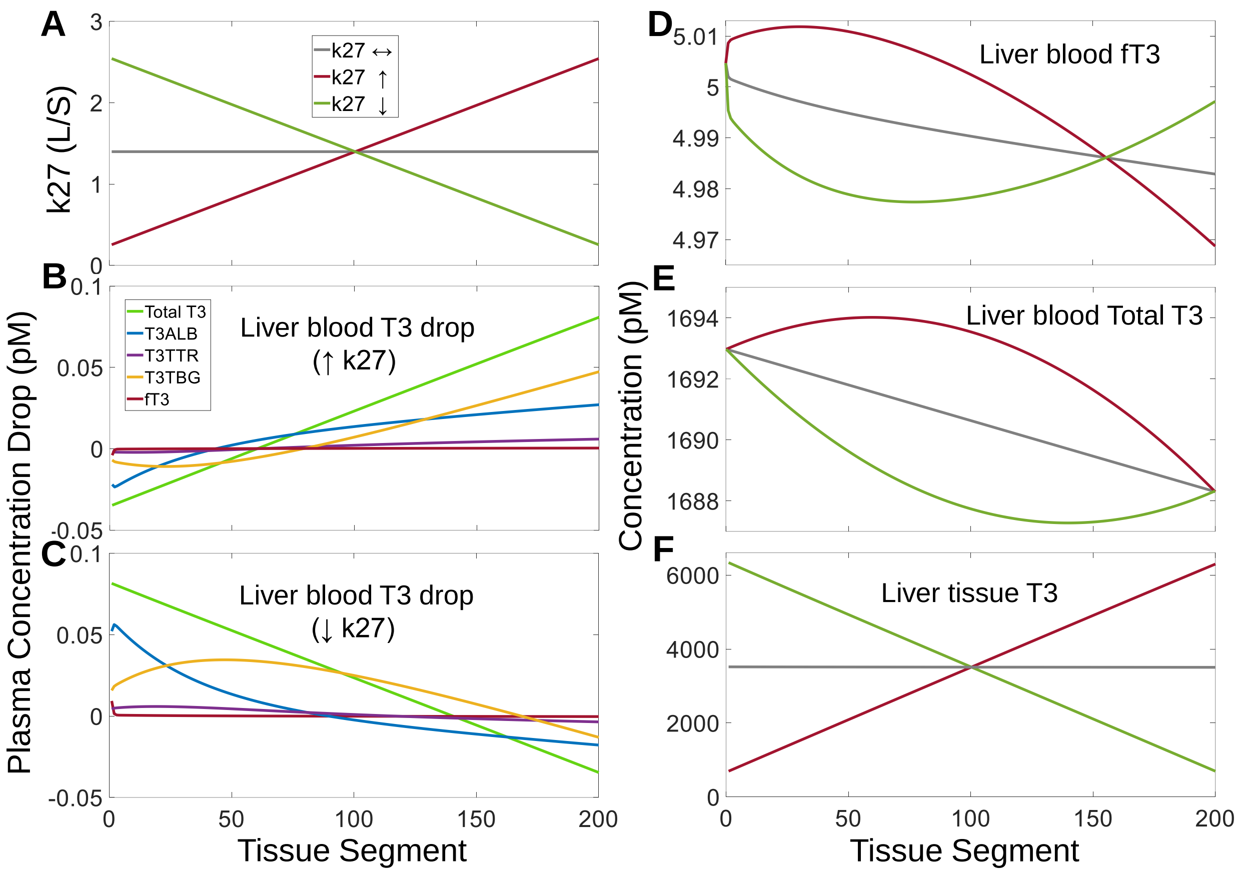


**Figure S8. Effects of $k_{27}$ gradients across *Liver* segments on T3 concentrations in *Liver* and contributions of THBPs to T3 loading and unloading in the spatial PBK model. (A)** Linearly increasing, decreasing, or constant $k_{27}$ gradients implemented as indicated. **(B-C)** Segment-to-segment net differences in plasma concentrations of *fT3*, *T3TBG*, *T3TTR*, *T3ALB*, and *Total T3* as indicated in *Liver blood* for increasing and decreasing $k_{27}$ gradients respectively. **(D-F)** T3 concentrations in *Liver blood* and *Liver tissue* as indicated. The same color scheme is used for panels (D-F) as indicated in (A), and for panel (C) as in (B).

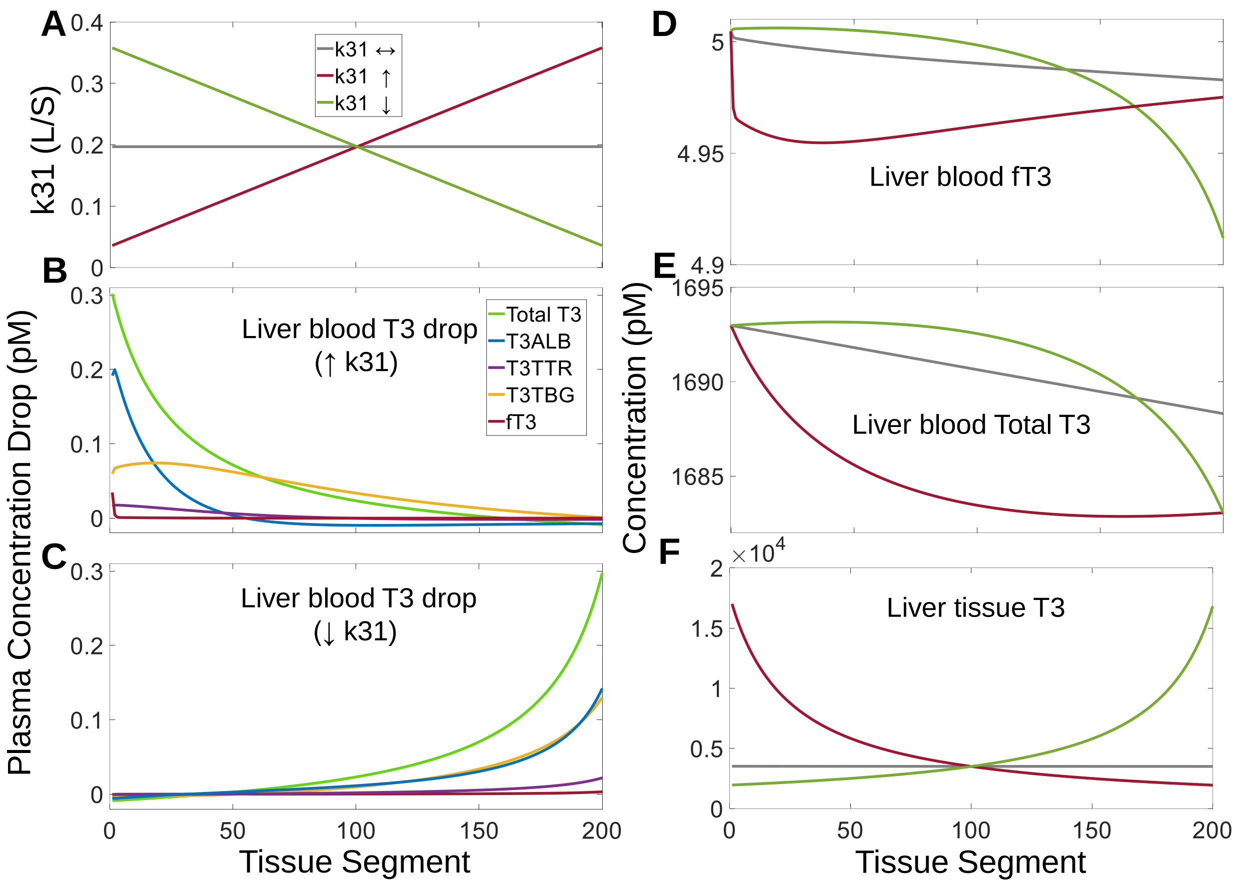


**Figure S9. Effects of $k_{31}$ gradients across *Liver* segments on T3 concentrations in *Liver* and contributions of THBPs to T3 loading and unloading in the spatial PBK model. (A)** Linearly increasing, decreasing, or constant $k_{31}$ gradients implemented as indicated. **(B-C)** Segment-to-segment net differences in plasma concentrations of *fT3*, *T3TBG*, *T3TTR*, *T3ALB*, and *Total T3* as indicated in *Liver blood* for increasing and decreasing $k_{31}$ gradients respectively. **(D-F)** T3 concentrations in *Liver blood* and *Liver tissue* as indicated. The same color scheme is used for panels (D-F) as indicated in (A), and for panel (C) as in (B). The nonlinearity of T3 in *Liver tissue* (F) and a greater drop in *Total T3* (E) levels can be likewise explained by the rationale described in the main text for Fig. 11 for the effects of $k_{30}$ on T4.

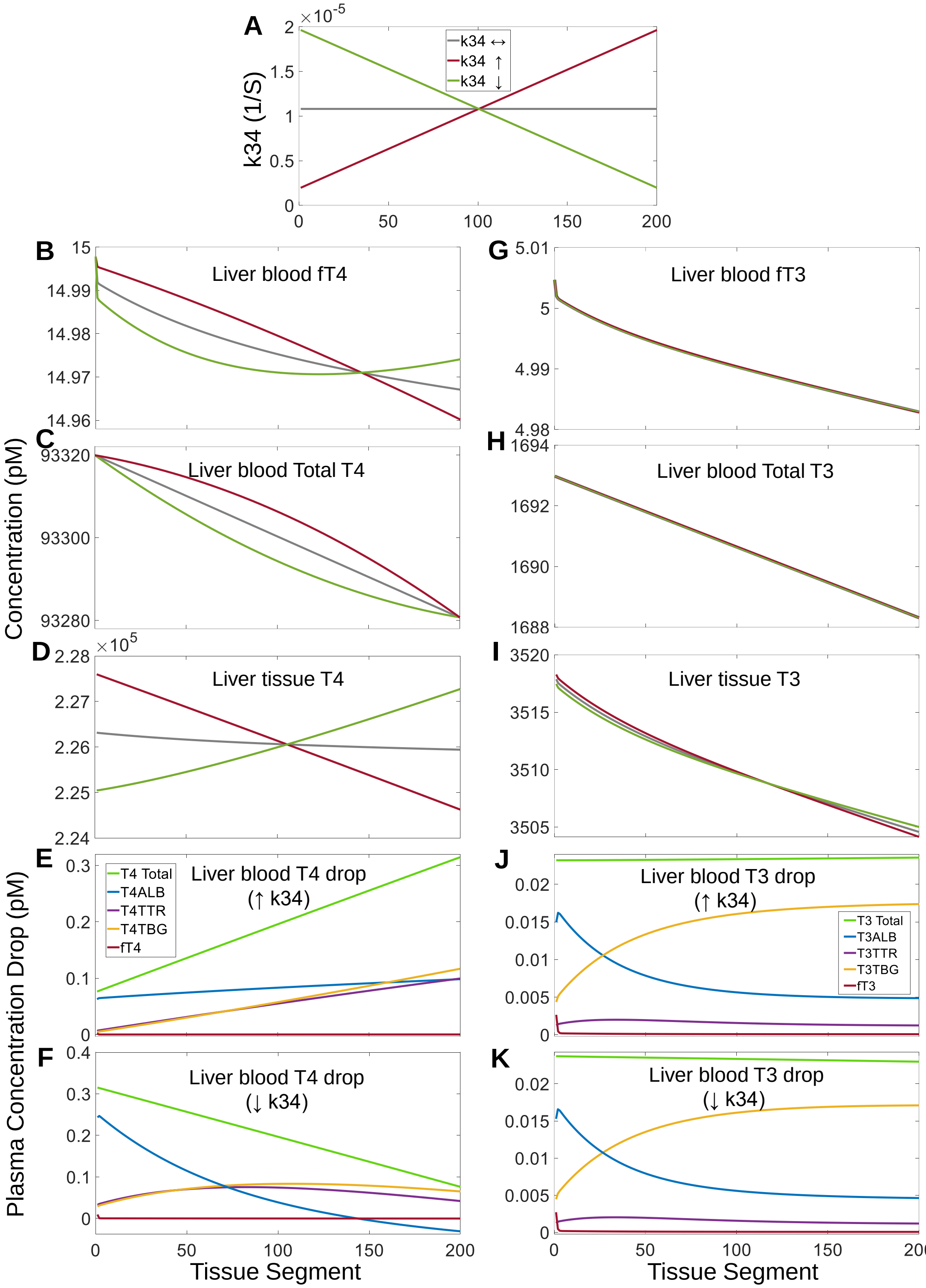


**Figure S10. Effects of $k_{34}$ gradients across *Liver* segments on TH concentrations in *Liver* and contributions of THBPs to TH loading and unloading in the spatial PBK model. (A)** Linearly increasing, decreasing, or constant $k_{34}$ gradients implemented as indicated. **(B-D)** T4

concentrations in *Liver blood* and *Liver tissue* as indicated. **(E-F)** Segment-to-segment net differences in plasma concentrations of *fT4*, *T4TBG*, *T4TTR*, *T4ALB*, and *Total T4* as indicated in *Liver blood* for increasing and decreasing $k_{34}$ gradients respectively. **(G-I)** T3 concentrations in *Liver blood* and *Liver tissue* as indicated. **(J-K)** Segment-to-segment net differences in plasma concentrations of *fT3*, *T3TBG*, *T3TTR*, *T3ALB*, and *Total T3* as indicated in *Liver blood* for increasing and decreasing $k_{34}$ gradients respectively. The same color scheme is used for panels (B-D, G-I) as indicated in (A), for panel (F) as in (E), and for panel (K) as in (J).

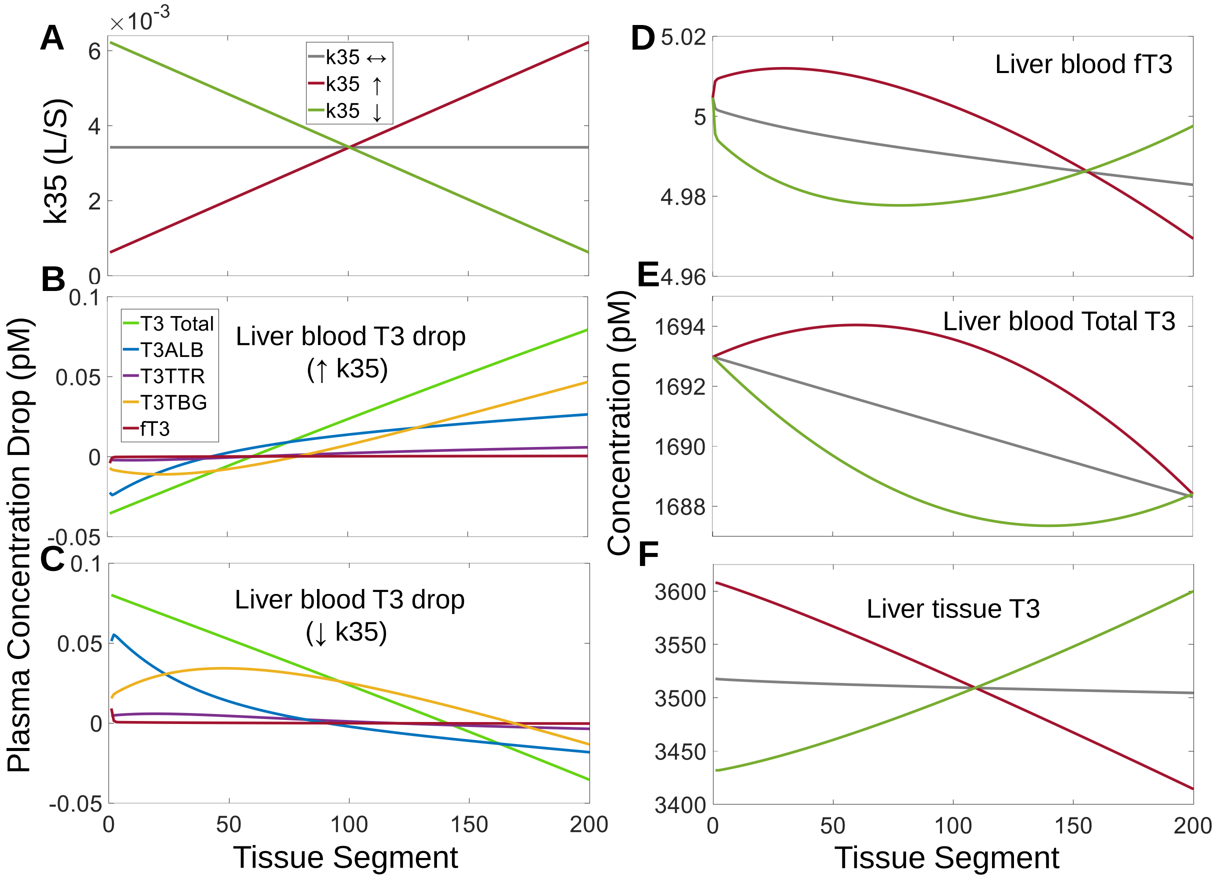


**Figure S11. Effects of $k_{35}$ gradients across *Liver* segments on T3 concentrations in *Liver* and contributions of THBPs to T3 loading and unloading in the spatial PBK model. (A)** Linearly increasing, decreasing, or constant $k_{35}$ gradients implemented as indicated. **(B-C)** Segment-to-segment net differences in plasma concentrations of *fT3*, *T3TBG*, *T3TTR*, *T3ALB*, and *Total T3* as indicated in *Liver blood* for increasing and decreasing $k_{35}$ gradients respectively. **(D-F)** T3 concentrations in *Liver blood* and *Liver tissue* as indicated. The same color scheme is used for panels (D-F) as indicated in (A), and for panel (C) as in (B).
