## Supplemental Tables for "Spatially Dependent Tissue Distribution of Thyroid Hormones by Plasma Thyroid Hormone Binding Proteins"

| **Table S1. Loading and unloading of THs in tissues by THBPs predicted by nonspatial PBK model** | | | | | | | | | | | | |
| --- | --- | --- | --- | --- | --- | --- | --- | --- | --- | --- | --- | --- |
|  |  | Free TH | |  | TH-TBG | |  | TH-TTR | |  | TH-ALB | |
|  |  | Conc  (pM) | CV-CA (pM) |  | Conc (pM) | CV-CA (pM) |  | Conc (pM) | CV-CA (pM) |  | Conc (pM) | CV-CA (pM) |
|  |  |  | % change |  |  | % change |  |  | % change |  |  | % change |
| T4 | CA | 15.0019 | na |  | 7.0046E4 | na |  | 1.6003E4 | na |  | 7.2754E3 | na |
|  | CV_Thyroid_ | 17.4036 | +2.4017 |  | 7.055E4 | +504 |  | 1.6467E4 | +463 |  | 8.1783E3 | +903 |
|  |  |  | +16.01% |  |  | +0.7198% |  |  | +2.895% |  |  | +12.41% |
|  | CV_Liver_ | 14.9745 | -0.0274 |  | 7.0031E4 | -15.175 |  | 1.5992E4 | -11.753 |  | 7.2631E3 | -12.264 |
|  |  |  | -0.1824% |  |  | -0.0217% |  |  | -0.0734% |  |  | -0.169% |
|  | CV_RB_ | 14.9901 | -0.0117 |  | 7.0034E4 | -11.782 |  | 1.5996E4 | -7.496 |  | 7.2698E3 | -5.592 |
|  |  |  | -0.0781% |  |  | -0.0168% |  |  | -0.0468% |  |  | -0.077% |
| T3 | CA | 5.0089 | na |  | 1.2758E3 | na |  | 82.2077 | na |  | 331.36 | na |
|  | CV_Thyroid_ | 5.9337 | +0.9247 |  | 1.3466E3 | +70.8 |  | 92.0188 | +9.811 |  | 383.59 | +52.23 |
|  |  |  | +18.46% |  |  | +5.55% |  |  | +11.93% |  |  | +15.76% |
|  | CV_Liver_ | 4.9879 | -0.021 |  | 1.2727E3 | -3.0267 |  | 81.9134 | -0.2943 |  | 330.04 | -1.3206 |
|  |  |  | -0.4194% |  |  | -0.237% |  |  | -0.358% |  |  | -0.3985% |
|  | CV_RB_ | 5.0045 | -0.0044 |  | 1.275E3 | -0.7841 |  | 82.1397 | -0.0681 |  | 331.07 | -0.2876 |
|  |  |  | -0.0879% |  |  | -0.0615% |  |  | -0.0828% |  |  | -0.08679% |
| Note: na, not applicable. CA: arterial concentration, CV: venous concentration, % change = (CV-CA)/CA. | | | | | | | | | | | | |

| **Table S2. Association and dissociation rates in Body Blood of spatial PBK model** | | | | |
| --- | --- | --- | --- | --- |
|  |  | **TBG** | **TTR** | **ALB** |
| **T4** | Association rate (pM/S) | 1260.682510329346 | 1331.150580835845 | 9455.265988830572 |
|  | Dissociation rate (pM/S) | 1260.590344780352 | 1331.101730743782 | 9455.406514398281 |
|  | Dissociation - association (pM/S) | -0.092165548993989 | -0.048850092063276 | 0.140525567709119 |
|  | % difference of association rate | -0.0073% | -0.0037% | 0.0015% |
| **T3** | Association rate (pM/S) | 210.2763628664731 | 56.657096966456798 | 728.1398981200394 |
|  | Dissociation rate (pM/S) | 210.2700894061734 | 56.657547088314935 | 728.1455890574169 |
|  | Dissociation - association (pM/S) | -0.006273460299610 | 0.0004501218581367539 | 0.0057 |
|  | % difference of association rate | -0.0030% | 0.00079% | 0.00078% |

| **Table S3. Association and dissociation rates in Thyroid blood of spatial PBK model** | | | | |
| --- | --- | --- | --- | --- |
|  |  | **TBG** | **TTR** | **ALB** |
| **T4** | Association rate (pM/S) | 1459.565671183635 | 1544.182645785973 | 10969.40610951351 |
|  | Dissociation rate (pM/S) | 1269.664970928976 | 1369.651802031955 | 10629.27496088456 |
|  | Dissociation - association (pM/S) | -189.9007002546589 | -174.5308437540182 | -340.1311486289433 |
|  | % difference of association rate | -13% | -11% | -3.1% |
| **T3** | Association rate (pM/S) | 248.6288214910533 | 67.122620306186676 | 862.7148940516059 |
|  | Dissociation rate (pM/S) | 221.9545034900577 | 63.427068997003730 | 843.0425469590634 |
|  | Dissociation - association (pM/S) | -26.674318000995527 | -3.695551309182946 | -19.672347092542509 |
|  | % difference of association rate | -11% | -5.5% | -2.3% |

| **Table S4. Association and dissociation rates in last segment of Liver blood (venous blood) of spatial PBK model** | | | | |
| --- | --- | --- | --- | --- |
|  |  | **TBG** | **TTR** | **ALB** |
| **T4** | Association rate (pM/S) | 1258.171572585189 | 1328.426386480677 | 9435.894730992155 |
|  | Dissociation rate (pM/S) | 1260.355005668355 | 1330.125087936212 | 9436.711779241517 |
|  | Dissociation - association (pM/S) | 2.183433083166619 | 1.698701455535002 | 0.817048249362415 |
|  | % difference of association rate | 0.17% | 0.13% | 0.0087% |
| **T3** | Association rate (pM/S) | 209.3727555530279 | 56.410532227945666 | 724.9695019198201 |
|  | Dissociation rate (pM/S) | 209.7857011151821 | 56.439659982927587 | 725.0831784258697 |
|  | Dissociation - association (pM/S) | 0.412945562154249 | 0.029127754981921 | 0.113676506049615 |
|  | % difference of association rate | 0.20% | 0.052% | 0.016% |

| **Table S5. Association and dissociation rates in last segment of RB blood (venous blood) of spatial PBK model** | | | | |
| --- | --- | --- | --- | --- |
|  |  | **TBG** | **TTR** | **ALB** |
| **T4** | Association rate (pM/S) | 1259.682105079881 | 1330.041598769491 | 9447.374499214060 |
|  | Dissociation rate (pM/S) | 1260.397025119185 | 1330.431177434313 | 9447.517914542790 |
|  | Dissociation - association (pM/S) | 0.714920039304161 | 0.389578664821556 | 0.143415328730043 |
|  | % difference of association rate | 0.057% | 0.029% | 0.0015% |
| **T3** | Association rate (pM/S) | 210.0990305580981 | 56.607075233117889 | 727.4959324422516 |
|  | Dissociation rate (pM/S) | 210.1413758741688 | 56.610011587762116 | 727.5077986501583 |
|  | Dissociation - association (pM/S) | 0.042345316070708 | 0.002936354644227 | 0.011866207906678 |
|  | % difference of association rate | 0.020 % | 0.0052% | 0.0016% |
